# Scaling concepts in ’omics: nuclear lamin-B scales with tumor growth and predicts poor prognosis, whereas fibrosis can be pro-survival

**DOI:** 10.1101/2021.02.25.432860

**Authors:** Manasvita Vashisth, Sangkyun Cho, Jerome Irianto, Yuntao Xia, Mai Wang, Brandon Hayes, Farshid Jafarpour, Rebecca Wells, Andrea Liu, Dennis E. Discher

## Abstract

Spatiotemporal relationships between genes expressed in tissues likely reflect physicochemical principles that range from stoichiometric interactions to co-organized fractals with characteristic scaling. For key structural factors within the nucleus and extracellular matrix (ECM), gene-gene power laws are found to be characteristic across several tumor types in The Cancer Genome Atlas (TCGA) and across single-cell RNA-seq data. The nuclear filament *LMNB1* scales with many tumor-elevated proliferation genes that predict poor survival in liver cancer, and cell line experiments show *LMNB1* regulates cancer cell cycle. Also high in the liver, lung, and breast tumors studied here are the main fibrosis-associated collagens, *COL1A1* and *COL1A2*, that scale stoichiometrically with each other and super-stoichiometrically with a pan-cancer fibrosis gene set. However, high fibrosis predicts prolonged survival of patients undergoing therapy and does not correlate with *LMNB1*. Single-cell RNA-seq data also reveal scaling consistent with the pan-cancer power laws obtained from bulk tissue, allowing new power law relations to be predicted. Lastly, although noisy data frustrate weak scaling, concepts such as stoichiometric scaling highlight a simple, internal consistency check to qualify expression data.

**Classification:** Applied Physical Sciences (major) and Cell Biology (minor)

**Significance Statement:** Non-linear scaling analyses pervade polymer physics and chemistry and conceivably provide new insight into polymeric assemblies of genes expressed in tissues as well as co-regulated gene sets. Fractal scaling and stoichiometric scaling are among the gene-gene power law results identified here for key structural polymers in nuclei or extracellular matrix in human cancer data. Among nuclear envelope factors that might scale with DNA mass, only one nuclear filament scales with tumor proliferation and predicts poor survival in some cancer types. Collagen-1 scales with fibrosis and also tends to increase in multiple tumor types, but patients in therapy surprisingly survive longest with the highest levels of fibrosis, consistent with a therapeutic response.

## Introduction

Tumors differ physically from normal tissues, often palpably. Breast tumor stiffness, for example, can be felt in self-exams and can affect tumor growth^1^. Fibrotic rigidity is a major risk factor for liver cancer^2^, and fibrosis is common in lung cancer, opposing lung compliance^3^. Stiffness changes are often attributed to abnormal tissue architecture and accumulation of structural proteins, especially collagens in extracellular matrix (ECM) that links via cell adhesions and cytoskeleton to nucleus. Dysmorphic nuclei also prevail in tumors, and sometimes reflect changes in nuclear lamins, which are primary determinants of nuclear shape and stiffness^4–7^. Lamins and collagens are among the most abundant architectural factors and can both affect cell cycle in model systems: lamins in cultured cells regulate senescence^8^ and telomere dynamics^9^, whereas proliferation is promoted by stiff ECM in 2D culture^10^ and 3D tumor models in mice^11^. Here, a physics-rooted analysis of human expression data across three major cancers – of liver, lung, and breast – link collagens and lamins to patient survival.

Collagens and lamins assemble into high-order fractals typical of structural polymers that impart architecture and properties^12^ – including stiffness scaling versus concentration^13^ with implications for gene-gene relationships (**Fig.1a**). We searched for power law scaling of collagens and lamins in relation to other genes quantified in The Cancer Genome Atlas (TCGA), and we complement the analyses with tumor proteomics data, cancer cell line cultures, and non-linear analyses of single-cell RNA-seq results. We look first in TCGA for coordinated changes in tumors relative to normal tissues in a given patient, and then for nonlinear gene-gene associations in tumors across patients – ultimately relating gene sets related by power laws to patient survival. Scaling laws in biology include Klieber’s well-known power law^14^ for increased metabolic rate versus (body mass)^3/4^. Although the molecular basis for such a law remains actively studied^15^, the scaling relations uncovered here pervade polymer physics and chemistry in terms of structures and properties (**Fig.1a**) or stoichiometry (**Fig.1b**). Such relations also underlie highly time-dependent biological processes such as cell cycle (**Fig.1c**).

**Fig.1.**
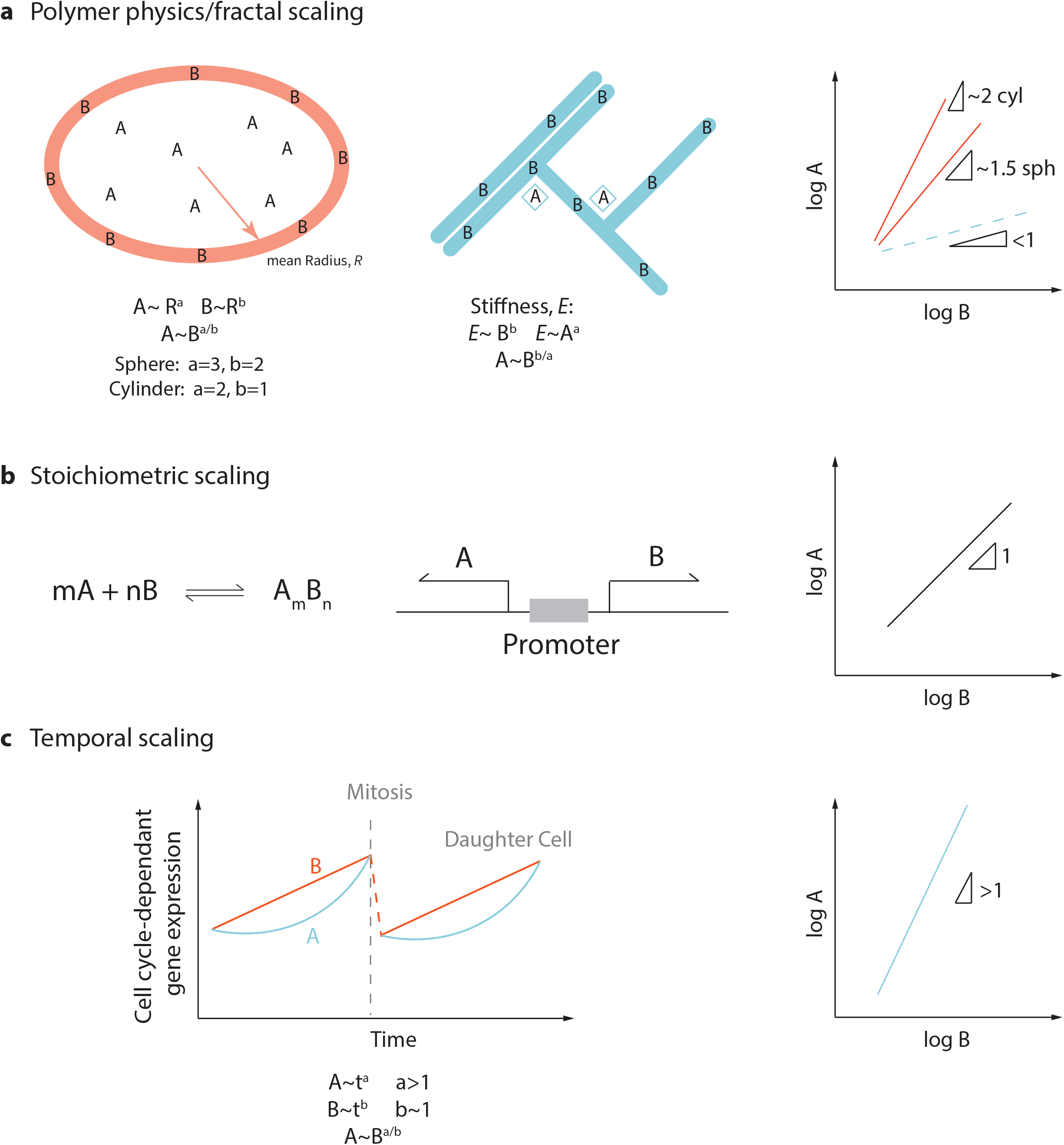
Physicochemical scaling concepts of expression. **a)** Factors A and B distribute differently and scale per fractal physics of a volume and its surrounding surface or according to branched, decorated networks. Mechanical properties that scale with such factors can also yield new scaling relations. **b)** Stoichiometric scaling is expected for co-regulated factors that form complexes or share a promoter. **c)** Genes that follow power laws in time can also scale with each other.

Collagen-I fibers are stoichiometric assemblies of COL1A1 and COL1A2 proteins, and a parallel increase of *COL1A1* and *COL1A2* transcripts (and protein) as a function of tissue stiffness has recently been shown useful in assessing data quality for normal tissues^16,17^. Such stoichiometric scaling is linear and can reflect co-regulated mRNA upstream of protein assemblies, including expression from the same promoter (**Fig.1b**). Paradoxically, while high collagen-I in fibrotic tissue associates with tissue stiffness and with tumorigenesis^1,2^, fibrotic reponses to therapy of several cancers in very small scale studies have also correlated with survival^18^. Changes in lamin levels and/or structure are likewise common in cancer^19^, but functional role(s) are unclear. Lamin-B1 (LMNB1) is normally the most invariant of the three lamin genes, but is elevated in liver tumors and is a circulating biomarker.^20,21^ *LMNB1* knockdown inhibits fibroblast proliferation, inducing senescence^8^, perhaps consistent with knockout mice dying at birth^22,23^, but *LMNB1’*s impact on cancer remains unclear^24^. Unlike LMNB1, lamin-A/C (*LMNA*) scales with tissue stiffness and collagen^25,26^ from soft brain to rigid bone^16^ and for some cancer lines^16,27^. Here, across three carcinomas, we identify sets of genes that scale with *LMNB1* and *separately* with collagen-I, and we relate the gene sets to surprisingly opposite relationships to patient survival.

## Results

### Expression scaling across tumors and normal tissues

Expression data in tumor-adjacent tissue (‘normal’) is provided for only a subset of patients in TCGA datasets, and so we analyze this data first from the perspective of collagen-I per:

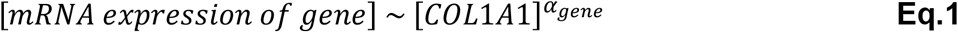

For *COL1A2* versus *COL1A1* (**Fig.2a**), expression in tumor is higher than that in adjacent tissue across liver, breast, and lung, consistent with a tendency for increased fibrosis in tumors^1,2^. Scaling exponents of α_COL1A2_ = 0.83 ± 0.11 (S.D.) are slightly lower than expected of stoichiometric scaling, but R^2^ > 0.84 likely indicates co-regulation over the large range of apparent expression (∼2^10^ fold). *COL4A2* versus *COL4A1* was examined for comparison because this main basement membrane collagen is expressed from the same promoter (**Fig.1b**) (unlike collagen-1), and scaling exponents are only ∼12% lower than stoichiometric scaling (R^2^ = 0.84 to 0.95) (**Fig.S1a**). Basement membrane *COL4A1* does not scale well with fibrous *COL1A1* (**Fig.S1b**), consistent with these ECM’s being distinct.

**Fig.2.**
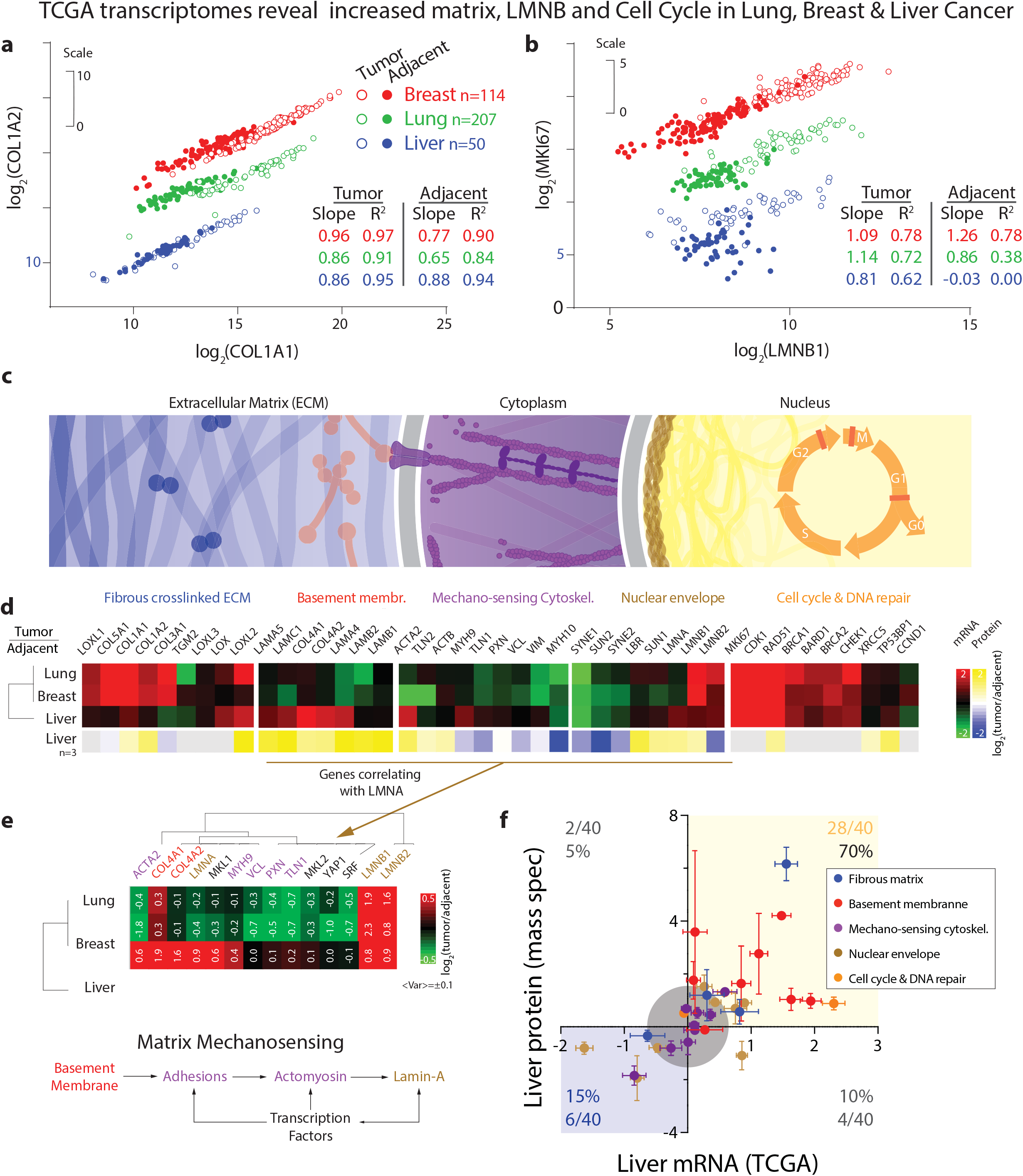
Fibrous ECM and filamentous Lamin-B’s increase with cell cycle in transcriptomes of tumors of Lung (n=207), Breast (n=114) and Liver (n=50) in The Cancer Genome Atlas (TCGA). **a)** RNA reads (RSEM) for the two subunits of the collagen-I heterotrimer scale together across all three cancers and corresponding adjacent tissue (R^2^ > 0.87). For each patient, tumor and adjacent tissue RNA are reported, and tumor shows more collagen-I on average. For clarity, Lung and Breast data are shifted up 3 and 6 units, respectively, and Lung Adenocarcinoma is the focus as the most common lung cancer^72^. **b)** RNA reads (RSEM) for proliferation marker *MKI67* versus nuclear lamina factor, *LMNB1*, reveals scaling in all three tumor types (R^2^ > 0.62) but not adjacent Lung or Liver tissue. For clarity, Lung and Breast data are shifted up 5 and 10 units, respectively, **c)** Schematic polymer systems of main interest. **d)** Standard red-green heatmap of log_2_(fold-change of RNA in tumor relative to adjacent tissue) in RSEM units averaged over all patients. Yellow-blue heatmap of log_2_(fold-change of protein in tumor relative to adjacent tissue) as obtained in LFQ units from mass spectrometry (MS) for n=3 Liver Cancer patients. **e)** RNA changes that correlate best with Lamin-A across the three cancers constitute a known matrix mechanosensitive pathway, but neither collagen-I nor lamin-B are in the pathway. **f)** Ratios of (tumor/adjacent) for Liver protein and RNA show 85% of data in the first and third quadrants.

Lamin-B1 filaments assemble constitutively around chromatin that of course increases in total mass with progression through the cell cycle, and so the power law

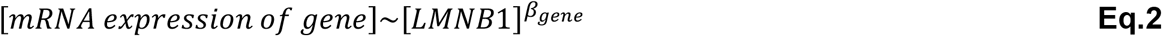

was first applied to the well-known proliferation marker *MKI67* (**Fig.2b**). Tumor data are again elevated relative to adjacent tissue, consistent with proliferation being a ‘hallmark’ of cancer, while the data also shows lung and breast (not liver) exhibit a reasonably continuous trend from normal to tumor over a ∼2^6^ range of expression (much narrower than collagen-1). All three tumor types show reasonable scaling with exponents from ∼0.8 to ∼1.3 (R^2^ = 0.6-0.8). However, normal liver and lung show no scaling over the narrowest ranges of *LMNB1*, suggesting low proliferation in normal tissue.

### Standard presentation: high Collagen-I, Lamin-B, & Cell Cycle, but varied Cytoskeleton-Lamin-A

Conventional presentation of gene trends shows the selected sets of structure-proliferation genes for lung and breast cancer (tumor/adjacent) data are similar but differ for liver cancer patients (**Fig.2c-f**). All three tumor types show similarly elevated levels of fibrillar collagens (collagens-1, 3, 5), cross-linking enzymes (lysyl-oxidases, LOX’s), B-type lamins (*LMNB1* and *LMNB2*), and cell cycle plus DNA-repair genes needed for progression through cell cycle checkpoints^28^.

Liver differs from lung and breast cancers (R^2^ = 0.4; **Fig.S1c**) in particular for basement membrane ECM (e.g. collagen-4), adhesions-cytoskeleton, and *LMNA*. Low Lamin-A/C in lung^29^ and breast^30^ cancers potentially relates to invasion based on studies of one lung cancer line that showed low *LMNA* favors 3D invasion *in vitro* and growth of tumors *in vivo*^31^. In various 2D cultures, LMNA drives adhesive spreading and actomyosin assembly^32^ downstream of the Serum Response Factor (SRF) pathway^27,33^, and high levels of *LMNA* in liver tumors (**Fig.2e**) and genes in the SRF pathway are indeed clear in hierarchical clustering with *SRF* and its co-activators (*MKL1, MKL2*). Similar trends are evident for *YAP1* in the Hippo pathway of growth^34^. *MKL1* associates with its well-established target genes, including adhesion factor vinculin (*VCL*)^35^ and nonmuscle myosin-IIA (*MYH9*)^27^ but also basement membrane collagen-4’s (unlike collagen-1). The results suggest basement membrane density, hence stiffness, is upstream of the known matrix mechanosensing adhesion-contractility-LMNA axis (**Fig.2e**). Importantly, *LMNB1* and *LMNB2* do not cluster within this mechanosensitive pathway.

Protein changes for liver (tumor/adjacent) conducted by quantitative Mass Spectrometry (MS) proteomics analyses (n=3 patients) show good concordance with TCGA transcriptome trends. Structural proteins are generally abundant, which is well-suited for MS proteomics. Overall, >85% of proteins and RNA showed concurrent upregulation/downregulation (**Fig.2f**). *LMNB1* was up in both analyses as were all three of the MS-detected DNA repair factors. Given the encouraging association between transcript and phenotype-dictating protein, we made use of the larger set of patients with tumor-only data (i.e. no adjacent sample) to ultimately relate TCGA expression to patient survival.

### Proliferative genes scale with *LMNB1* and predict poor survival

Whole genome analysis of transcripts in primary tumor tissue from liver cancer patients (**Fig.3a-c**, n=371) shows *LMNB1* expression varies by the same 32-fold range as the smaller dataset with normal tissue (**Fig.2b**, n=50). Scaling exponents *β*_*gene*_ and R^2^ of the fits for all 17,958 genes (per Eqs.1,2) are illustrated for proliferation-related genes *TOP2A, CDK1* and *MKI67* (**Fig.3a,b, S2a**). *TOP2A* is among the top-24 most upregulated genes (tumor versus adjacent normal) together with fifteen other genes (many associate with proliferation) that also scale well with *LMNB1* (**Table S1**). An anti-correlation for the hepatocyte-expressed, complement-related factor *MASP2* (**Fig.3c**) is consistent with a shift away from differentiation given that such genes accumlate in a protracted cell cycle^36^. Anti-correlations differ in lung cancer (**Fig.S2b**), consistent with a distinct lineage.

**Fig.3.**
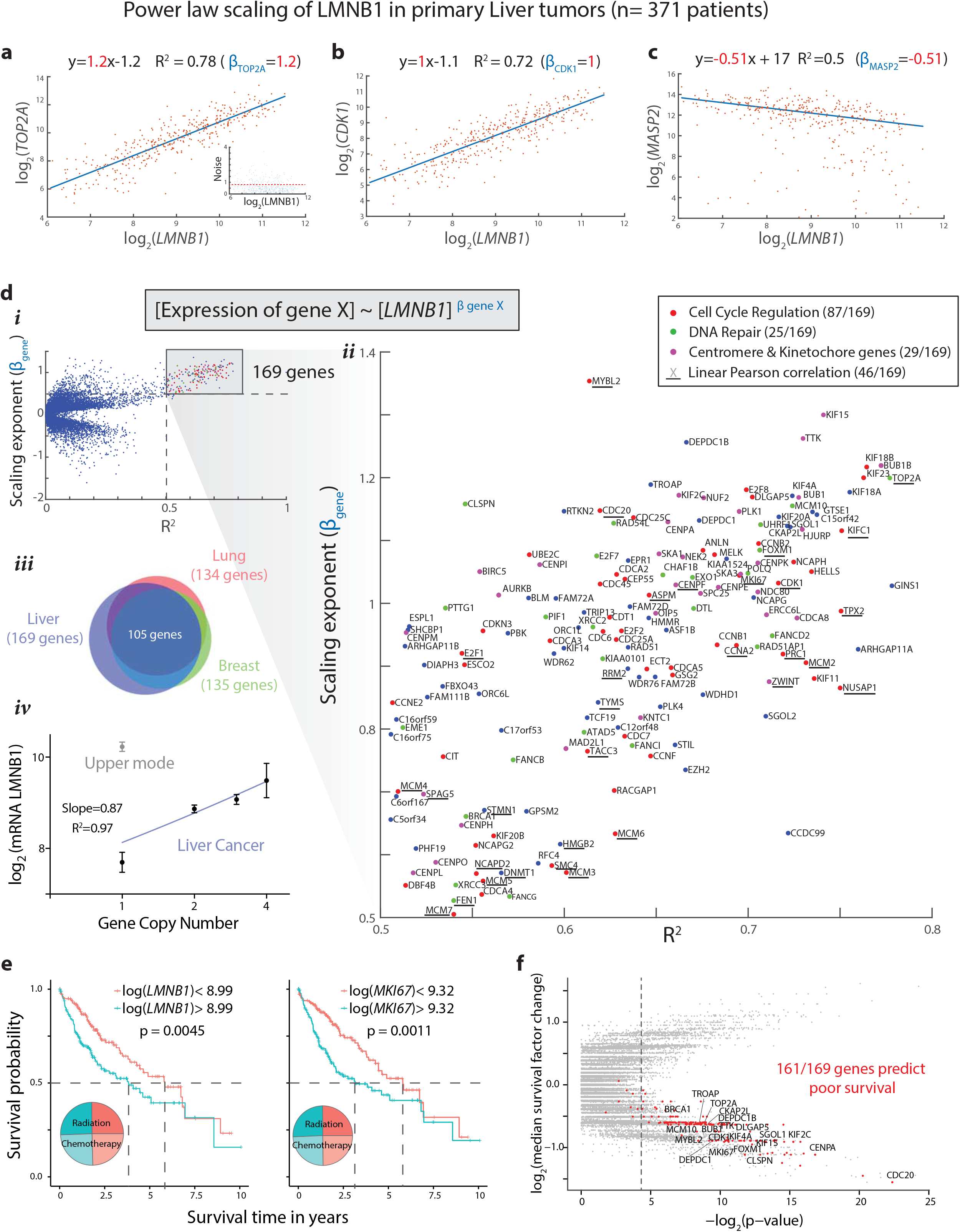
Power law scaling of *LMNB1* in Liver tumors (n=371 patients) **a-c)** Plots versus *LMNB1* RNA for correlated cell cycle genes *TOP2A* and *CDK1* as well as for anti-correlated gene *MASP2*. The R^2^ and scaling exponent (*β*_*gene*_) are indicated. **d)** Genes that correlate with *LMNB1* RNA. **i)** For all 17,958 genes, *β*_*gene*_ versus R^2^ gives a sideways-volcano plot. **ii)** 169 genes scale strongly *β*_*gene*_> 0.5 and well R^2^ > 0.5, and most relate to cell cycle regulation. Only a small subset of genes show significant Pearson correlations (fit on linear-linear plot; underlined). **iii)** Venn diagram shows the numbers of genes with *β*_*gene*_ > 0.5 and R^2^ > 0.5 in Lung (n=515 patients) and Breast Cancer (n=1,097 patients); 105 genes are common between the three datasets. **iv)** Gene copy numbers of *LMNB1* in liver cancer patients is almost linear in RNA expression, although patients with a single copy are bimodally distributed and only the lower mode fits the trend. **e)** For patients with *LMNB1* or *MKI67* RNA levels that exceed the respective median levels for all patients, the median time for survival is significantly shorter (p < 0.05) by 2-3 yrs in Kaplan Meier (KM) plots. **f)** For all 17,958 genes, similar KM analyses are summarized by the fold-change in median survival plotted against the p-value, yielding 3,464 genes that show significant differences, including 161 (of 169) genes that scale with *LMNB1* and show poor survival when expression exceeds the median.

Fitted parameters for all genes yield a ‘sideways-volcano plot’ (**Fig.3d-*i***). Most well-correlated genes (i.e. *β*_*gene*_>0.5, R^2^>0.5) are proliferation-related (83% of 169 genes: **Fig.3d-*ii*,*iii***). Fits on linear scales (i.e. Pearson) yield a small fraction of these genes (underlined in plot), but scaling with respect to proliferation marker *MKI67* yields 153 well-fit genes, >95% of which are identical to the *LMNB1* gene set (**Fig.S2c**) – which underscores invertibility. To assess whether *LMNB1* transcript levels relate to the number of copies of the *LMNB1* gene, genomic data for each liver cancer patient was analyzed^37^, noting that copy number variations are consistent with malignancy and can predict patient survival^38–41^. Transcript levels binned on gene copy number fit a power law of ∼1 (**Fig.3d-*iv***) – although patients with one copy of the *LMNB1* gene are bimodal distributed and only the low-expressors fit the trend. Consistent with gene dosage DNA → mRNA → protein, our proteomics for liver cancer detected LMNB1 plus three proliferation factors that are all upregulated in tumors (**Fig.2b,e**).

Survival data for liver cancer (371 patients) shows median survival is only ∼3-4 yrs for patients with high *LMNB1* or high *MKI67*, whereas low expressors survive almost twice as long (**Fig.3e**). This partitioning is independent of the fact that all patients are treated with either radiation or chemotherapy (**Fig.3e, inset pie-chart**), and independent of etiology including alcohol or hepatitis (**Fig.S2d**). Across all genes, the median survival for high/low expressors shows significant Kaplan-Meier plots for 3,464 genes, and among the genes (2,111) that predict poor survival when expression is high are >95% of the genes that scale with *LMNB1* (**Fig.3f**). Other genes are pro-survival, including high *MASP2* (**Fig.S2e**), consistent with a *LMNB1* anti-correlation (**Fig.3c**). Staging of primary tumors in terms of size and invasion also shows *LMNB1* increasing (**Fig.S2f**), whereas *LMNA* and *COL1A1* show no trends. A sideways-volcano plot for *LMNA* shows only one gene that scales strongly (**Fig.S2g**), which underscores the distinctive significance of *LMNB1*.

Sideways-volcano plots of *LMNB1* for lung and breast cancer (**Fig.S3a,b**) yield an overlapping set of the same genes: 78% of such genes in lung tumors (515 patients) are identical to those in breast (1,097 patients) and liver cancer (**Fig.3d-*iii***). Such pan-cancer scaling suggests coordinated regulation of proliferation, and strong scaling (exponent β > 1.1) hints at (Volume/Area) scaling (**Fig.1-i**). Regardless of mechanisms, Kaplan-Meier plots for *LMNB1*-scaling genes in primary lung tumors include *LMNB2, BRCA1, MKI67*, and *TOP2A* (**Fig.S3a, S4a,b,c**), all of which show poor survival for high expression (**Fig.S4d,e**). Recent studies of proliferating, embryonic cardiomyocytes suggest *LMNB2* facilitates attachment of microtubules to centromeres^42^. Such survival predictions are perhaps consistent with *LMNB1*-scaling in tumor but not normal liver tissue (per **Fig.2b**) and with *LMNB1* being a marker of proliferation.

### Perturbing *LMNB1* levels perturbs cancer cell cycle

Several approaches were used to assess LMNB1’s potential regulatory role in cell cycle. First, *LMNB1* was gene-edited with red fluorescent protein (RFP) and likewise for GFP-histone-H2B, using the endogenous promoters to drive expression in the lung cancer cell line A549^43^ and avoiding potential artifacts of overexpression. Live-cell imaging shows RFP-LMNB1 intensity per nucleus increasing as a power law in time up to mitosis (per **Fig.1c**), at which point the lamina reversibly disassembles^44^, and then re-assembles in daughter cells at half the original intensity (**Fig.4a,b**). Importantly, *A* ∼ *B*^α/β^ scaling with cell cycle at the single cell level also applies to the same genes (see **Theory Supplement**) in tumors with differences in mean doubling time – consistent with scaling between cancer patients (**Fig.3a-d**).

In fixed cells, RFP-LMNB1 scales with DNA-staining intensity across ‘2N’, ‘4N’, and higher ploidy cells (**Fig.4c**), which agrees with the very similar scaling with time of RFP-LMNB1 and GFP-histone-H2B (**Fig.4b**).Such results are confirmed with U2OS osteosarcoma cells and immunostaining of lamin-B (**Fig.4d**). The simplest spherical model of a nucleus with LMNB1 ∼ (Area of the nucleus)^1^ and with fractal filling by DNA ∼ (Volume of the nucleus)^δ^, gives DNA ∼ LMNB1^1/ δ^, so that cell culture suggests δ ∼ 1. In comparison, the scaling exponent of the top dozen scaling transcripts for proliferation and replication relative to *LMNB1* is β ∼ 0.9-1.3 (**Fig.3d**). Furthermore, DNA ∼ LMNB1 is robust against *LMNA* knockdown (Fig 4c), consistent with the lack of scaling of *LMNA* with *LMNB1* (**Fig.3d**), and with the distinct roles for the two lamin genes in mechanosensing (*LMNA*) (**Fig.2f**) and proliferation (*LMNB*’s).

**Fig.4.**
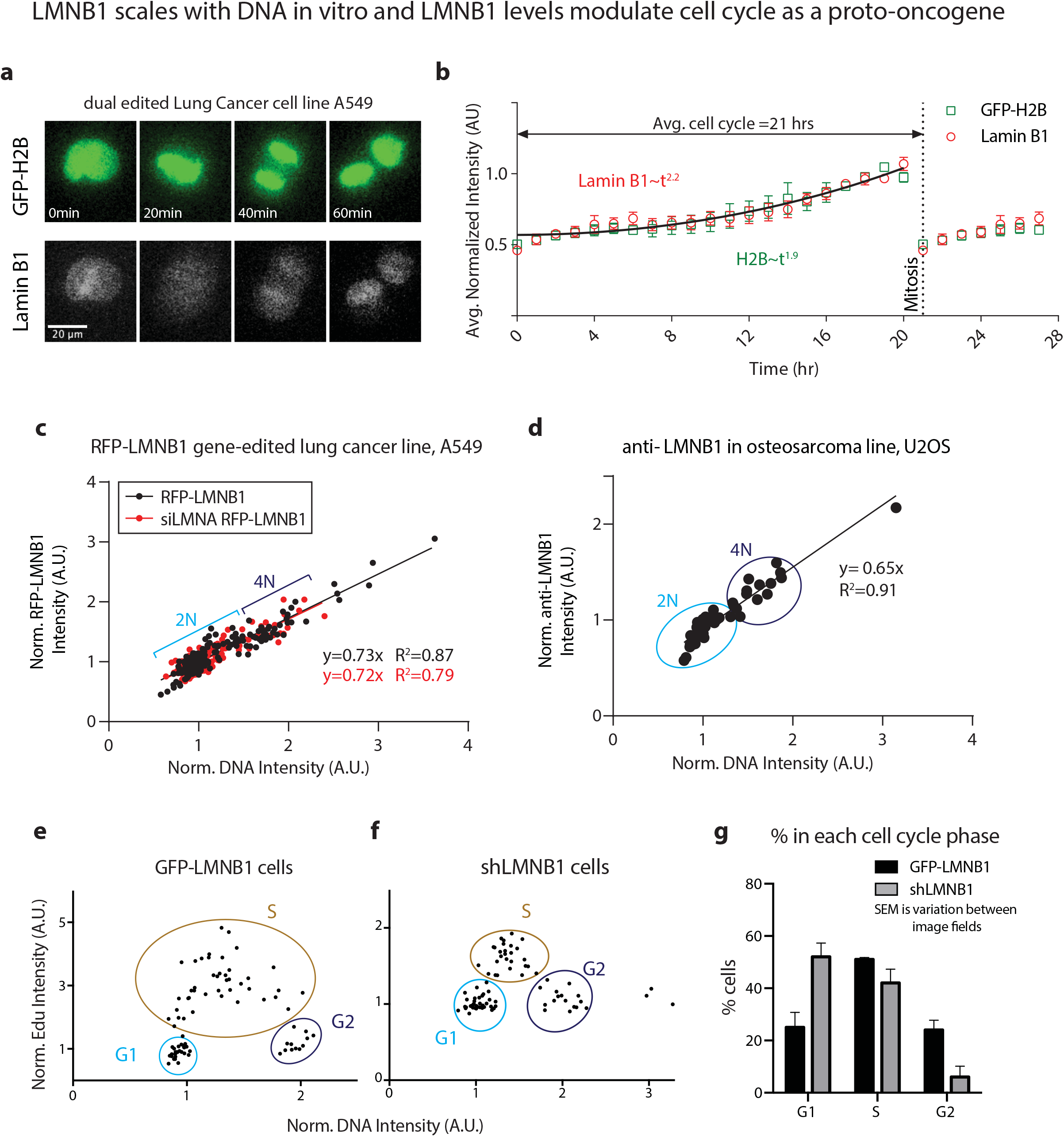
LMNB1 scales with DNA *in vitro* and LMNB1 levels modulate cell cycle as a proto-oncogene, **a,b)** Live cell imaging of gene-edited lung cancer line expressing GFP-H2B and RFP-LMNB1 shows parallel increases in intensities normalized to telophase in mitosis, with a mean cell cycle of 21±0.35 hrs. **c)** Intensity of RFP-LMNB1 is linear versus Hoechst-stained DNA in fixed A549s, even with lamin-A knockdown. Intensities are normalized to cells in the non-replicated state, ‘2N’. anti-LMNB also shows LMNB increases linearly with DNA in osteosarcoma-derived U2OS cells. **e-g)** EdU incorporation (1 hr) in replicating cells in combination with Hoechst-stained DNA identifies cell cycle stage (G1/S/G2). Knock down with shLMNB1 in U2Os cells reveals a smaller percentage of cells proceeding to S and G2 compared to overexpressing GFP-LMNB1 cells. Error bars indicate SEM values across image fields

To assess effects of LMNB1 on cancer cell proliferation, we transfected GFP-LMNB1 plasmid or shLMNB1 plasmid into U2OS cells, and pulsed incorporation of the nucleotide analog EdU (for 1h) was combined with DNA staining to quantify cell cycle stage^43^. High EdU signal indicates ongoing DNA synthesis (S-phase), and low EdU signal indicates G1 or G2 phases, depending respectively on low or high DNA intensity, i.e. ‘2N’ or ‘4N’ (**Fig.4e**). Knockdown cells show more cells in G1 compared to wildtype or overexpressing cells (identified by GFP signal at the single-cell level), whereas the latter were more in S-phase and G2 phase (**Fig.4f**), implying change in cell cycle progression with *LMNB1* changes (**Fig.4g**). These cancer line results agree with fibroblast results for which knockdown arrests cell cycle^8^, and they suggest LMNB1 has a proto-oncogene effect.

### ECM fibrosis genes scale with *COL1A1* across cancers and predict liver cancer survival

Both collagen-1 and lamin-B1 are high in all three cancers (**Fig.2a,b,d**), and so scaling with *COL1A1* was examined in a second whole genome analysis of TCGA transcripts, starting with primary tumors of liver cancer patients (n=371). *COL1A1* expression varies ∼16,000-fold (versus ∼32-fold for LMNB1 in **Fig.5a**), and does not relate to genome copy number changes in *COL1A1* (**Fig.S5a**), consistent with expression by non-malignant stromal cells. Furthermore, even though fibrous COL1A1 and associated ECM are abnormally high within a given patient across cancers (**Fig.2d**), between patient tumors these genes do not correlate with *LMNB1* (**Fig.5a**: α_LMNB1_ ∼0 and R^2^ < 0.1). However, *COL1A2* again scales almost linearly with *COL1A1*: α_COL1A2_ = 0.86 (**Fig.5b**); in comparison, an ECM protease gives α_MMP2_ = 0.79 (**Fig.5c**). The *COL1A2* scaling also has the highest R^2^ among all 17,958 genes, and the only other transcripts with R^2^ > 0.9 are *COL5A1* and *COL3A1* (**Fig.5d**), which are fibrous collagens known to associate with collagen-1 fibers^45^; both also scale with COL1A1 in proteomics^16^. Among 162 genes that scale with *COL1A1* (α_gene_ > 0.5, R^2^ > 0.5), the majority are ECM genes (**Fig.5d-*i*,*ii***), but also notable are *PI3K/AKT* signaling and *GLI2* (16 genes). Additional genes fit well to weak scaling (α_gene_ < 0.5, R^2^ > 0.5), and include basement membrane collagens *COL4A1* and *COL4A2*, suggestive of a (COL1 / COL4) ∼ (bulk/surface) scaling (**Fig.1-i**).

Across lung, breast, and liver cancers, 29 ‘fibrotic genes’ consistently scale strongly with *COL1A1* (**Fig.5d-*iii***; **Fig.S5b,c**), and most are ECM genes. One exception is *FAP*, fibroblast activating protein, which is a membrane-bound gelatinase targeted in the clinic^46^. Missing is the widely-analyzed, fibrosis marker in the cytoskeleton *ACTA2*^47^ (α-smooth muscle actin), and an analysis of genes that scale with *ACTA2* across all three cancers yielded 17 pan-cancer ‘myo-fibroblast genes’ (**Fig.5d-*iv***), which included cytoskeletal factors known to co-express with *ACTA2* (e.g. *CNN1*) but no ECM genes. Just one factor is shared with the 29 ‘fibrotic genes’ (*AEBP1*). Restricting the comparison to liver cancer, 9 genes scale with both *COL1A1* and *ACTA2* (**Fig.S6a**).

**Fig.5.**
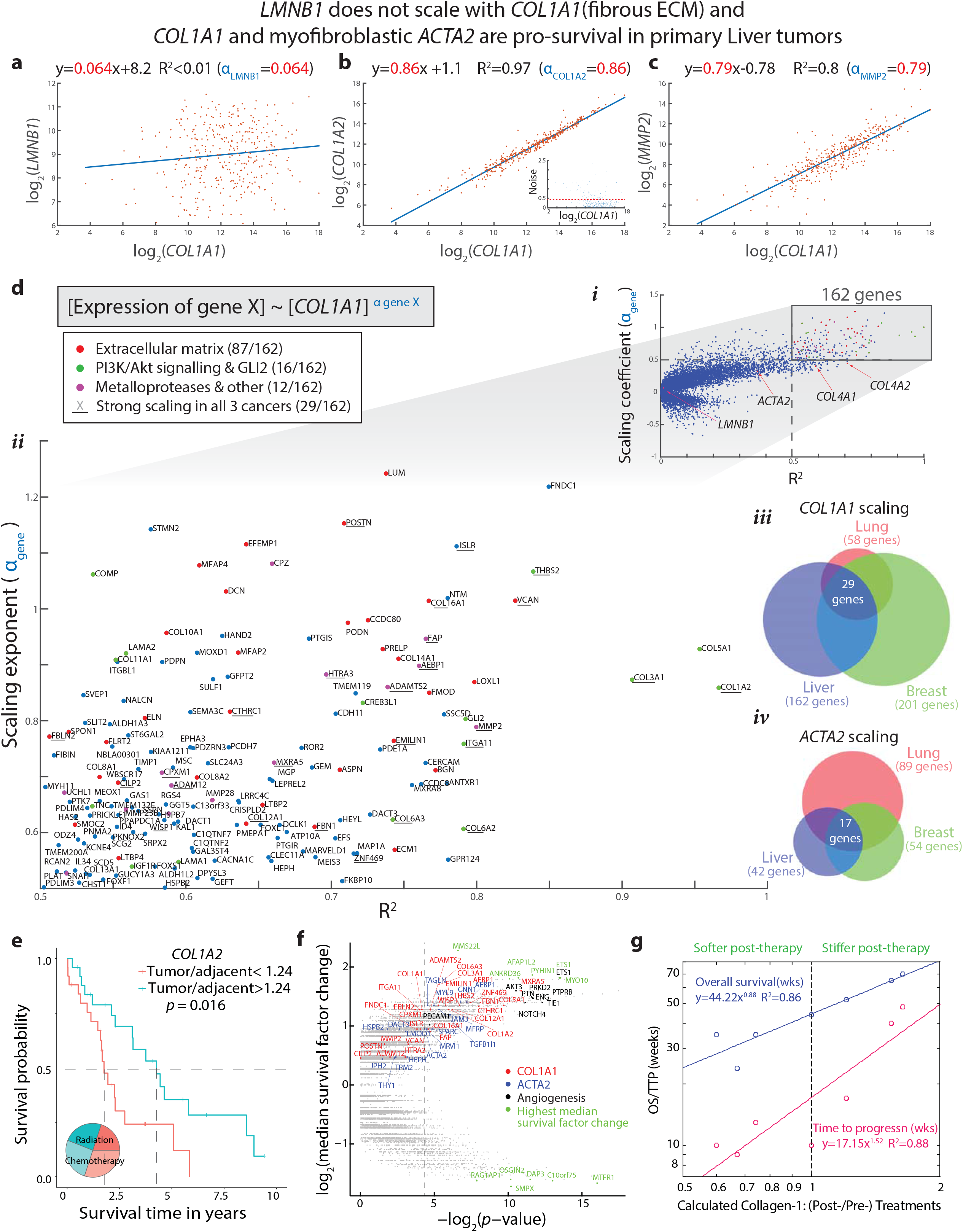
*LMNB1* does not scale with *COL1A1* (fibrous ECM) and *COL1A1* and myofibroblastic *ACTA2* are pro-survival in primary Liver tumors (n=371 patients) **a-c)** Plots versus *COL1A1* RNA for *LMNB1* (not correlated) and correlated ECM genes *COL1A2* and *MMP2*. The R^2^ and scaling exponent (*a*_*gene*_) are indicated. **d)** Genes that correlate with *COL1A1* RNA. **i)** For all 17,958 genes, *a*_*gene*_ versus R^2^ gives a sideways-volcano plot. **ii)** 162 genes scale strongly *a*_*gene*_ > 0.5 and well R^2^ > 0.5, and more than half relate to ECM. **iii)** Venn diagram shows the numbers of genes with *a*_*gene*_ > 0.5 and R^2^ > 0.5 in Lung (n=515 patients) and Breast Cancer (n=1,097 patients); 29 genes are common between the three datasets. **iv)** Similar Venn diagram for myofibroblastic gene *ACTA2* shows 17 genes in common. **e)** For *COL1A2* RNA levels (tumor/adjacent) that exceed the median levels for all patients (n=50 patients), the median time for survival is significantly longer (p < 0.05) by 2-3 yrs. **f)** For all 17,958 genes, KM analyses are summarized by the fold-change in median survival plotted against the p-value, showing that many of the genes which scale with *COL1A1* or with *ACTA2* predict prolonged survival when the expression ratio (tumor/adjacent) exceeds the median. Select angiogenesis and extrema genes are also indicated. **g)** Liver tumor stiffness measured before therapy and 6 wks later^18^ was converted to calculated changes in collagen-1 (Col1 ∼ E^1.5^) and plotted versus patient survival (n=7 patients), revealing strong power laws.

Kaplan-Meier plots for the subset of patients with patient-specific results for (tumor/normal) gene expression show *prolonged* survival for *COL1* (∼2 yrs versus ∼4yrs in **Fig.5e**) and for *ACTA2* (**Fig.S6b**); indeed, *all* 29 pan-cancer ‘fibrotic genes’ and *all* 17 pan-cancer ‘myo-fibroblast genes’ in a whole genome analysis of liver cancer are pro-survival (**Fig.5f**). Kaplan-Meier analyses of patients with tumor-only measurements (n=371) are compelling for genes that scale with *ACTA2* (**Fig.S6c-f**). Different etiologies such as alcohol or hepatitis also show no impact on survival predictions (**Fig.S6g**), and the partitioning is again independent of the fact that all patients are treated with either radiation or chemotherapy (**Fig.5e, inset pie-charts**).

*Pro-survival* results for *high* expression of fibrotic and myofibroblast genes in treated liver cancer patients appears consistent with results from a recent study of patients with liver stiffness measured by magnetic resonance elastography (MRE) before immunotherapy and 6 wks afterwards.^18^ Remarkably, patient survival correlated positively with a post-treatment *increase* in local liver stiffness (rather than initial or final stiffness). Such results for stiffness (*E*) are readily converted to collagen-I changes through the relation COL1A1 ∼ *E*^1.5^ obtained from mass spectrometry of tumors and diverse normal tissues^12^. Power law fits for Time-to-Progression and Overall Survival (in weeks) both reveal strong scaling over a ∼2^2^ fold calculated range of collagen-I that increases or decreases after treatment (**Fig.5g**).

### Pan-cancer power laws for genes scaling with *COL1A1, ACTA2*, or *LMNB1*

Genes that scale well with *COL1A1* for all three cancers in TCGA show similar pan-cancer power laws (**Fig.6a**), as does *COL4A2* versus *COL4A1* (**Fig.6b**). The latter pan-cancer exponent of 0.95 ± 0.02 indicates stoichiometric scaling (**Fig.1**). Mass spectrometry derived scaling results^12^ included not only COL1A1 ∼ *E*^1.5^ but also COL5A1 ∼ *E*^1.3^, which yields COL5A1 ∼ COL1A1^0.87^. TCGA’s pan-cancer scaling *COL5A1* ∼ *COL1A1*^0.82-0.92^ agrees as do exponents for other genes. Such agreement of tumor and normal transcript and protein is evidence of ‘universality’ in scaling.

**Fig.6.**
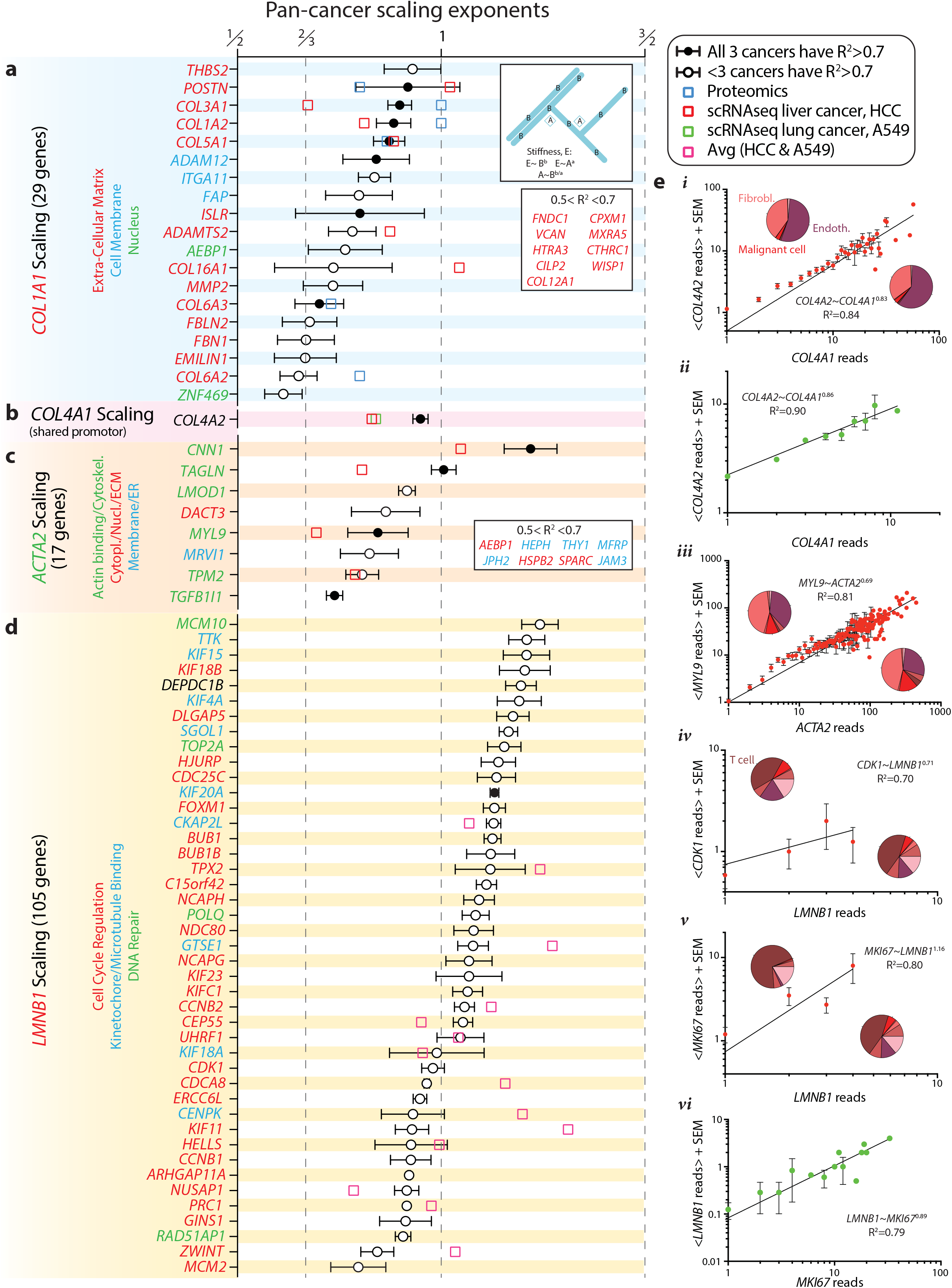
Pan-cancer power law exponents. **a-d)** (circle) Exponents for genes in TCGA that show indicated scaling with slope > 0.5 and R^2^ > 0.5 across Liver, Lung and Breast cancer. (blue square) Exponents obtained from proteomics across normal tissues and model tumors. Exponents from single-cell RNA-seq: (red square) hepatocellular carcinoma, HCC; (green square) lung cancer derived A549 line; (pink square) average exponent of HCC and A549. **e)** Power law fits of raw reads from single-cell mRNA sequencing. **i-ii)** *COL4A1* vs *COL4A2* from HCC patients or A549 cells. Pie charts for HCC: <5% of malignant cells have non-zero reads for *COL4A1* or *COL4A2*, versus ∼60% of Endothelial Cells, ∼35% are Fibroblasts. **iii)** *ACTA2* vs *MYL9* from HCC scale together as myofibroblastic genes and show similar cell-type specific expression profiles. **iv-vi)** *LMNB1* vs *CDK1* or *MKI67* for HCC or A549’s. For HCC, ∼45% of cells with non-zero reads of *LMNB1* or *CDK1* are Tcells, and <5% are malignant cells.

All exponents for *COL1A1* are also <1, suggesting collagen-1 fibers branch throughout the volume of a tumor or tissue, whereas the other factors occupy a fraction of branches (**Fig.1a**). Consistent with such a fractal scaling also for the cytoskeleton, genes that scale reproducibly with *ACTA2* exhibit scaling exponents as high as 1.2 and down to 0.6 across cancers (**Fig.6c**). A similarly broad range of exponents (0.6 to 1.3) for *LMNB1* (**Fig.6d**) suggests higher values (>1) are consistent with a lamina that surounds replicating DNA (**Fig.1a**) or perhaps a bias from early or late cell cycle genes (**Fig.1c**). Bulk sequencing of tumors is done, afterall, on a diversity of cell types at different cell cycle stages, which motivates a final examination of power laws in single-cell RNA-seq (scRNA-seq).

### Scaling in single-cell RNA-seq

Hepatocellular carcinoma (HCC) tumor biopsies from chemotherapy treated patients have only recently been characterized by scRNA-seq, which provided data from diverse cell types^48^. Not all cell types are equally extractable from a stiff solid tissue as intact single cells, which necisitates bulk analysis as above, but scRNA-seq certainly provides some important insights. Tumor fibroblasts and endothelial cells express *COL1A1, COL4* isoforms (**Fig.6e-*i***: exponent = 0.83), and *ACTA2* – all with up to ∼100 reads or more in scRNA-seq, while malignant HCC cells do not express these genes. Stoichiometric scaling of *COL4A1* versus *COL4A2* was nonetheless confirmed with lung cancer A549 cells (**Fig.6e-*i***: exponent = 0.86), and *COL4* scaling in TCGA adjacent and tumor (**Fig.S1a**: exponent = 0.88 ± 0.10) and the larger TCGA data (exponent = 0.95) show power law exponents from scRNA-seq generally correspond to those from bulk TCGA for pan-cancer exponents (**Fig.6a-e**). The scRNA-seq results, all based on raw reads, otherwise emphasize power law scaling over the usual linear, Pearson-type analyses.

Insights into cellular composition of tumors are possible with scRNA-seq. For example, tumors that are in high *ACTA2* in TCGA’s bulk sequencing (**Fig.5d-*iv***) potentially indicate more fibroblasts and endothelial cells based on scRNA-seq (**Fig 6e-*iii***). However, *LMNB1* in HCC tumors is detected moreso in easily extracted (non-adherent) Tcells than malignant cells and other cell types (**Fig 6e-*iv,v*; Fig.S7a,b**) even though A549s clearly express *LMNB1* (**Fig.6e-*iv*; Fig.S7c**). Nonetheless, scaling exponents for *LMNB1-*scaling genes that were averaged for HCC and A549 scRNA-seq again correspond closely to those from TCGA’s pan-cancer exponents (**Fig.6d**). Deviations likely result from low reads from *LMNB1* and other mitotic genes: only 8 out of the 105 pan-cancer genes have >10 reads per cell, including *MKI67*. This scRNA-seq analysis is reasonably consistent with ‘universality’ in scaling.

## Discussion

Across liver, lung, and breast carcinomas, specific sets of genes scale together with *LMNB1* in proliferation and separately with *COL1A1* in fibrosis, and at least for liver cancer, the distinct gene sets produce highly divergent predictions for patient survival. Metastasi of liver carcinoma (HCC) is less common than lung and breast carcinomas, and lamin-B1 is not only elevated in malignant hepatocytes but is even a circulating biomarker^20,21^. Relationships to malignant cell proliferation have been unclear but are understandable even though signatures for some tumors might be dominated by proliferation of immune or stromal cells that respond to patient treatments (**Fig 6e-*i,iii-v***). LMNB1 is constitutively expressed in all cells starting with invertebrates^49,50^ and associates closely with chromatin post-mitosis^51,52^, whereas lamin-A is highly mechanosensitive and is easily detected only as embryonic tissues stiffen^9^. Lamin-B1 knockdown undermines early development unlike *LMNA*^53^, and cell culture studies show *LMNB1* depletion inhibits DNA replication^54-55^ – all of which is consistent with cell cycle scaling with *LMNB1* and not *LMNA* (**Fig.3d, S3a-b, S2h**). Such a difference is also evident in now-standard analyses of scRNA-seq data with dimensional reduction and clustering (**Fig.S7a**), which confirms actively proliferating cells colocalize high *LMNB1* with *MKI67* and *TOP2A*, whereas *LMNA* is more broadly expressed across cell types and more similar to *COL4’s* (**Fig S7b,c**).

Upregulation of fibrous ECM is common in solid tumors (e.g. **Fig.2a,d**) and is surprisingly pro-survival in liver cancer – at least in therapy (**Fig.5f,g**). *COL1* subunit transcripts scale nearly stoichiometrically with each other and scale with other fibrous collagens and ECM genes (e.g., *COL3, COL5, LOXL1*; **Fig.5d**), adding confidence to the conclusion that collagen-I changes are independent of LMNB1. Because *COL1A1* power laws for the pan-cancer fibrosis set are all <1, it increases super-stoichiometrically with respect to the other genes; a reasonable analogy is that collagen-1 forms the woody bulk of a tree that the other factors assemble onto like bark or leaves on the tree. Further insight is provided by scRNA-seq of HCC, with high expression of *COL4’s* and *ACTA2* in endothelial and fibroblasts whereas *COL1’s* are only in fibroblasts (**Fig.S7b**). Nonetheless, both fibrous *COL1A1* and basement membrane *COL4A1* scale not only with other liver ECM genes but also with *PI3K*/*Akt* and the *GLI2* hedgehog pathway (**Fig.5d, S8a**) that causes scarring in persistent activation^56^.

Distinct from both *LMNB1* and *COL1* is a >20-fold increase in *COL15A1* in tumor vs normal tissue (**Table S1**). Such an increase has been attributed elsewhere to vascular myofibroblasts that enhance vacularization in fibrosis^57^. Increased *COL15A1* is consistent with scaling (**Fig.S8b**) and with vascular genes such as *TIE1* that prolong survival of treated liver cancer patients (**Fig.5f,S8c-h**). One study that also identified pro-survival roles for fibrosis and myofibroblasts in a mouse model of pancreatic cancer also described a pro-survival role for vascular genes^58^. The most enriched Gene Ontology (GO) annotations in survival are indeed extracellular factors, exosomes, and metabolic pathways (**Fig.S6c**), with examples of the latter genes including high levels of P450 and alcohol dehydrogenase genes that suggest maintenance or regeneration of a well-differentiatied liver ^59,60,61–64^.

Pro-survival fibrosis and vasculogenesis signatures of cancer patients in therapy (**Fig.5g**) could result from multiple mechanisms. High vascularization will improve efficacy of therapy, for example, and as cancer cells die, a fibrotic in-filling is likely as an *acute* wound-healing response. Increased tumor stiffness after therapy^65^ would indeed be consistent with a so-called ‘pathological response’ to pre-surgical chemoradiation, as is known in multiple cancers including pancreatic cancer to favor survival^65^. Non-invasive measurements of stiffness (MRE) and perfusion (various radiological methods) might help distinguish effects of *acute* stiffening from *chronic* stiffening which is a clear risk factor for cancer^66–68^.

### Scaling is interchangable, but weak scaling is lost in noisy TCGA data

Conventional heatmap presentations can suggest associations, such as between *COL1A1* and *LMNB1* (**Fig.2c-f**), where none might exist. However, a key strength of quantitative power law relationships is the ease of predicting new power law relations:

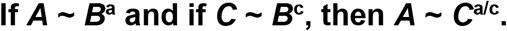

For example, defining genes *A* = *COL5A1, B* = *COL1A1*, and *C* = *COL1A2* with measured power laws of *a* = 0.86, *c* = 0.93 (**Fig.6a**), gives *a/c* = 1.08. Measurements agree: *COL5A1* ∼ *COL1A2*^1.06^ (R^2^ = 0.96). Poorer fits yield poorer predictions of course, but weak scaling genes are also generally problematic.

Genes scale weakly or strongly with *LMNB1* and *COL1A1* across the three cancers in sideways-volcano plots (i.e. R^2^ > 0.5) are rare or non-existent. This is surprising given evidence of genes that scale weakly (α < 0.5) with *COL1A1* in small datasets^17^, but a likely reason is illustrated by the noise in *COL1A2* scaling (**Fig.5b, inset**: RMSE = 0.44) – which we apply to generic power law functions *f* that are weaker (or stronger) power laws versus *COL1A1* than the actual data. Power law fits of these equally noisy *f* yield R^2^ values that decrease strongly from high R^2^ with the best-fit exponents, with the same trends in *LMNB1* analyses (**Fig.S9**). Noisy data thus explains the lack of weak scaling genes in TCGA data, and noise in the liver cancer data likely reflects batch effects (i.e. 19 batches of patient samples analyzed over ∼3 yrs in several sequencing centers). Nonetheless, the strong scaling genes and especially the ‘universal’ power laws supported by proteomics-rheology and initial scRNA-seq (**Fig.6**) should motivate new physicochemical theories for the interacting pathways that underlie proliferation and ECM production as well as patient survival.

## Methods & Materials

### TCGA Analysis

Gene expression (mRNA-seq), copy number variation (Gistic 2 thresholded) and phenotype data was downloaded from UCSC Xena website (https://xenabrowser.net/datapages/). Primary tumor sample can be identified by the code 01, for example, TCGA-69-7978-**01** is and TCGA-69-7978-**11** is solid tissue normal or “uninvolved” adjacent tissue from the same patient. In the Liver Cancer data set, the histological type includes 6 patients with Hepatocholangiocarcinoma (one set of normal and tumor tissue both), 3 patients with Fibrolamellar Carcinoma and 362 Hepatocellular Carcinoma patients. 32.4% of the patients in the primary tumor Liver Cancer dataset are female. If >75% of the patients showed 0 reads for a gene, i.e. out of 371 primary tumor samples in Liver Cancer, if >278 patients showed 0 reads, that gene was removed from analysis. This reduced the Illumina HiSeq gene expression dataset from 20,530 genes to 17,958 genes. For cancer staging, TCGA annotated TNM classification system was used, which pathologically stratifies the primary tumors (T) from T1-4 indicating increasing tumor size. Amount of spread into nearby structures was chosen to characterize the primary tumor size phenotype T1 (Solitary tumor without vascular invasion), T2 (Solitary tumor with vascular invasion or multiple tumors, none > 5 cm), T3 (Multiple tumors > 5 cm; or Macrovascular invasion), T4 (Tumor(s) with direct invasion of adjacent organs or with visceral peritoneum).

### Statistical Analysis

Kaplan Meier plots were created using the libraries survminer and survival in R version 3.3.3. Survival fits using survfit function were made using survival object(surv), which contained time and event (death or left the study) data. The Kaplan Meier plot and p-value were obtained using the ggsurvplot function. Box-plots were made using the function boxplot in R. One-way ANOVA test (which tests significant differences in the averages of multiple data groups) was conducted using aov function in R. Linear fit models for log values of gene expression were made using the fitlm function in MATLAB 2019b using the robust regression model that iteratively reweights least squares to reduce the effect of outliers, making it less sensitive than ordinary least squares to large changes in small parts of the data. For example, sideways-volcano plots made with robust regression model for *LMNB1* in primary Liver tumors give 169 genes with high β_gene_ and R^2^, but with ordinary linear regression, there are 159 genes.

R^2^ measures the improvement in prediction of the variable ‘y’ (eg, log_2_(mRNA expression of *TOP2A*)) using the regression model *f*, based on predictor x (eg, log_2_(mRNA expressionof *LMNB1*)), compared to the ‘mean model’ (**Eq 3**) (i.e. does the variation in ‘x’ predict variation in ‘y’ better than just using the mean value of ‘y’ as the predictor).

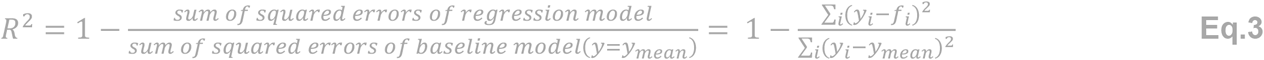

### Human liver collection

Human liver samples were collected in strict accordance with the protocols approved by the Institutional Review Board of the University of Pennsylvania and the Philadelphia Veterans Administration Medical Center (protocol number 820896). De-identified samples for this research study were obtained from male and female patients aged 32 to 68 years old who underwent liver resection or transplantation. Liver tissue was initially stored in cold Hank’s Balanced Salt Solution (HBSS, Gibco), then flash frozen until analysis by mass spectrometry.

### Mass spectrometry (LC-MS/MS) of Liver tumors and adjacent uninvolved tissues

Tissues were lysed using NUPAGE LDS buffer. The lysates were loaded onto SDS-PAGE gels (4-12%, Bis-Tris, Invitogen) and gel electrophoresis was run for 20 min at 160 V. ∼1 mm^3^ gel sections were carefully excised from SDS–PAGE gels and were washed in 50% 0.2 M ammonium bicarbonate (AB), 50% acetonitrile (ACN) solution for 30 min at 37°C. The washed slices were lyophilized for >15 min, incubated with a reducing agent (20 mM TCEP in 25 mM AB solution), and alkylated (40 mM iodoacetamide (IAM) in 25 mM AB solution). The gel sections were lyophilized again before in-gel trypsinization (20 mg/mL sequencing grade modified trypsin, Promega) overnight at 37°C with gentle shaking. The resulting tryptic peptides were extracted by adding 50% digest dilution buffer (60 mM AB solution with 3% formic acid) and injected into a high-pressure liquid chromatography (HPLC) system coupled to a hybrid LTQ-Orbitrap XL mass spectrometer (Thermo Fisher Scientific) via a nano-electrospray ion source.

Raw data from each MS sample was processed using MaxQuant (version 1.5.3.8, Max Planck Institute of Biochemistry). MaxQuant’s built-in Label-Free Quantification (LFQ) algorithm was employed with full tryptic digestion and up to 2 missed cleavage sites. The software’s decoy search mode was set as ‘revert’ and a MS/MS tolerance limit of 20 ppm was used, along with a false discovery rate (FDR) of 1%. The minimum number of amino acid residues per tryptic peptide was set to 7, and MaxQuant’s ‘match between runs’ feature was used for transfer of peak identifications across samples. All other parameters were run under default settings. The output tables from MaxQuant were fed into its bioinformatics suite, Perseus (version 1.5.2.4), for protein annotation and sorting.

### Cell lines

A549 (human lung adenocarcinoma) cell lines were obtained from ATCC (American Type Culture Collection, Manassas, VA, USA) and U2OS cell lines were obtained from the laboratory of Roger Greenberg, University of Pennsylvania. ATCC provides cell line authentication test recommendations per Tech Bulletin number 8 (TB-0111-00-02; 2010). These include: microscopy-based cell morphology check, growth curve analysis, and mycoplasma detection (by DNA staining) which were conducted on all cell lines used in these studies. All cell lines maintained the expected morphology and standard growth rates with no mycoplasma detected. U2OS and A549 cell lines were cultured in DMEM high-glucose media and Ham’s F12 nutrient mixture (Gibco, Life Technologies), respectively, supplemented with 10% fetal bovine serum (FBS) and 1% penicillin and streptomycin (Sigma-Aldrich). shLMNB1 retrovirus construct was obtained from the laboratory of Prof. Robert Goldman (Northwestern University, Chicago), prepared as previous described^8^. GFP-LMNB1 plasmid was obtained from the laboratory of Profs. Harald Herrmann-Lerdon and Tatjana Wedig. Cells were transfected using Lipofectamine 2000 (1:1000, Thermo fisher scientific) and incubated for 3 days.

### Gene editing and live cell imaging analysis

CRISPR/Cas9 and homology directed repair were used to incorporate a full length GFP tag sequence in the genomic loci for H2B at the C-terminus, using mEGFP (K206A). We used the ribonucleic protein (RNP) method with recombinant wild type S. pyogenes Cas9 protein pre-complexed with a synthetic CRISPR RNA (crRNA) and a trans-activating crRNA (tracrRNA) duplex. Both the crRNA and the tracrRNA were purchased from Dharmacon, while the Cas9 protein was purchased from UC Berkeley QB3 Macrolab. The sequence for the crRNA is 5’ ACTCACTGTTTACTTAGCGC 3’, and for the tracrRNA is 5’ GTCGCAGTCGCCATGGCGGG 3’.

The crRNA:tracrRNA duplex was prepared my adding crRNA (20 uM) and tracrRNA (20 uM) at a 1:1 ratio for a final concentration of 10 uM. The mixture was heated at 95°C for 5 min and then allowed to cool at room temperature for 2 hrs. After 2 hrs elapsed, 700,000 A549 RFP-LMNB1 cells were prepared and suspended in 200 uL of Ham’s F12 media supplemented only with 10% FBS. Cells were placed in a 0.4 cm Gene Pulser electroporation cuvette (Bio-Rad). 10 uL of 10 uM crRNA:tracrRNA duplex, 10 uL of 10 uM Cas9 protein, and to the cell suspension was also added 8 ug of donor plasmid (AICSDP-52: HIST1H2BJ-mEGFP is Addgene plasmid # 109121). Cells were then electroporated using a Gene Pulser Xcell electroporation system at 160 V for 30 ms. The RFP-LMNB1 expressing A549 cells were as described elsewhere, with RFP tag insertion done using zinc finger nuclease at the C-terminus. The RFP tag is a monomeric TagRFP from Evrogen.

Successfully edited cells were enriched using fluorescence-activated cell sorting (FACS) with a BD FACSJazz (BD Biosciences). Cells were enriched three times using FACS. Single-cell clones were then prepared by plating 100 cells into a 15-cm plate (Corning) and isolated using cloning rings (Corning). To confirm GFP tagging of H2B by PCR, the forward primer sequence is 5’ ATGCCAGAGCCAGCGAA 3’, and the reverse primer sequence is 5’ GAGAGTTTGCAACCAACTCACT 3’. Successful H2B integration, then a PCR product equivalent to the sum of the GFP and H2B molecular weights is detected in addition to endogenous H2B. To confirm RFP insertion, PCR analysis was performed. RFP integration was confirmed using the RFP forward primer (sequence 5’ AAGAGACCTACGTCGAGCAGC 3’) and a LMNB1 reverse primer (sequence 5’ AAAGCGGGAGCTCCTTCTCGGCAGT 3’). A PCR product was only detected for RFP-tagged cells with this primer set.

For imaging, A549 RFP-LMNB1 GFP-H2B were freshly sorted for double positive cells and seeded in a 24 well plate. Fresh media was replaced the next day, and cells were imaged at 20X magnification for 36 hrs with 20 image fields taken per well using the EVOS FL Auto imaging system (Thermo Fischer Scientific) at the CDB Microscopy Core at the Perelman School of Medicine.

### Immunofluorescence imaging

Cells were rinsed with pre-warmed PBS on a shaker at low speed for 3 min, fixed with 4% paraformaldehyde (PFA, Fisher) for 15 min, washed 3 times with PBS and permeabilized with 0.5% Triton-X (Fisher) in PBS for 30 min. Permeabilized cells were then blocked with 5% BSA in PBS for ≥1.5 hrs. Samples were incubated overnight with primary antibodies in 0.5% BSA solution, with gentle agitation at 4 °C. Hoechst 33342 (1:1000, Thermo Fisher, #3570) was used to stain the DNA. The primary antibodies used were: LMNA (1:500, Cell Signaling Technology (CST), #4777), LMNB (1:500, Santa Cruz, #sc-6217). Samples were washed three times in 0.1% BSA in PBS and incubated with the corresponding secondary antibodies at 1:500 dilution for 1.5 hrs at RT (Alexa Fluor 488 and 647 nm; Invitrogen). Cells on glass coverslips were mounted with mounting media (Invitrogen ProLong Gold Antifade Reagent). Images of adherent cells were taken with an Olympus IX81. All images in a given experiment were taken under the same imaging conditions and analyzed using ImageJ (NIH).

### EdU labeling and staining

EdU (10 µM, Abcam) was added to 24 well plates 1 hr before fixation. Cells were fixed and permeabilized as described above. After permeabilization, samples were stained with 100 mM Tris (pH 8.5) (MilliporeSigma), 1 mM CuSO4 (MilliporeSigma), 100 µM Cy5 azide dye (Cyandye), and 100 mM ascorbic acid (Millipore Sigma) for 30 min at room temperature. Samples were washed three times with 15-min washes on the shaker and then underwent immunostaining as described above.

### Single cell RNA-seq and data analysis

A549 cells were seeded in a 96 well-plate at a density of 150,000 cells/well and cultured for 2 days. Cells were then trypsinized and submitted to the Center for Applied Genomics (1014, Abramson Research Center, University of Pennsylvania) for single-cell RNA sequencing (with the 10X Genomics single-cell gene expression kit, a 400M read flow cell & single-end read using HiSeq from Illumina). 2,453 cells were sequenced with an average of 64,404 reads (and 3,457 genes) per cell. Sequencing saturation is only 49.4% and 97.5% of reads can be mapped to genome. Data were extracted from R using Cell Ranger R kit and aligned to genes via BioMart. Normalization to transcript length is not required because 10X Genomics only sequences the 3’ end.

For all cells that express a particular integer value, for example 4 reads of *COL4A1*, we determine for *COL4A2* (gene *y*) the average number of non-integer reads = <*y*>. Plotting these in a log-log plot (excluding *x* = 0 and <*y*> = 0 reads), we determined the best-fit scaling exponent, and then repeated for *COL4A2* (gene *x*) and *COL4A1* (gene *y*) to obtain the inverse exponent. Error in y indicates S.E.M. COL4A1 and COL4A2 log-log slope fits have an inverse correlation and their averages converge to values close to 1.

Public GEO database code: GSE125449 was used in the Single cell sequencing analysis from hepatocellular carcinoma patients. The following patient codes were used in the analysis: S02_P01_LCP21 (704 cells), S07_P02_LCP28 (124 cells), S10_P05_LCP23 (151 cells), S12_P07_LCP30 (805 cells), S15_P09_LCP38 (1046 cells), S16_P10_LCP18 (124 cells), S21_P13_LCP32 (132 cells)

For average single-cell exponents with respect to *LMNB1*, only genes showing R^2^ > 0.7 were included. Identification of cell type based on sequencing data was performed as previously described^48^.

### Visualization of Dimensionally reduced Single cell mRNA-seq

Seurat^69^ was used to analyze Single cell sequencing data of A549 cells and HCC tumor cells which are mentioned above. For HCC tumor dataset, cells with >50% mitochondrial genes were removed since they were probably dead cells. Cell >6000 unique gene reads were removed as those are the droplets with doublets/multiplets of cells in each droplet. Cells are then normalized by total expression in each cell and then multiplied by 10000 and log transformed. Reads were then scaled, followed by principal component analysis of the variable features. This was followed by clustering analysis with resolution 0.1. Elbow Plot was used to determine that the first 10 principle components capture 90% of the variation. Uniform Manifold Approximation and Projection (UMAP) analysis was done to dimensionally reduce the first 10 principle components, followed by visualization. Similar analysis was done for the A549 cells.

## Author Contributions

Conceptualization, MV, DED; Investigation, MV, SC, YX, MW, BH, DED; Validation, MV, DED; Formal Analysis, MV, DED; Resources, RW; Writing, MV, DED; Supervision, DED; Funding Acquisition, DED

## SUPPLEMENTAL THEORY

### Population Scaling from Single Cell Scaling

Consider quantities *A* and *B* that scale with the age (time since last division) *a* of cells. Then the population values of *A* and *B* are given by the averages *A(a)* and *B(a)* over the distribution *ρ*(*a*)of ages *a* in the population. In a growing population with the doubling time *τ*, the age distribution is approximately given by (see derivation below)

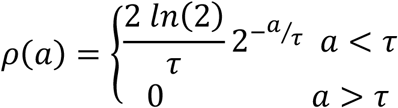

For *A*(*a*)∝ *a*^*α*^ and *B*(*a*)∝ *a*^*β*^, we have

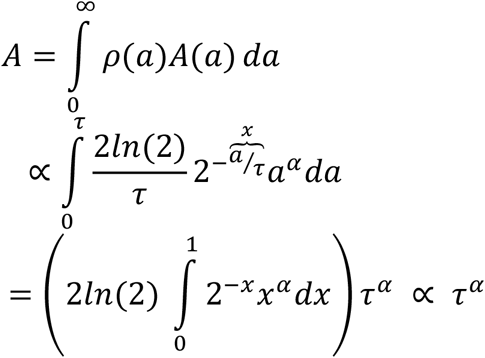

and similarly, *B* ∝ *τ*^*β*^. If *A* and *B* are sampled across cancer patients with some variability in the doubling time *τ* across the patients, *B* would scale with *A* as 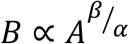.

While this argument only includes the variability in population doubling time *τ* across population samples, the resulting scaling is robust with respect to small heterogeneities in cell cycle times within each populations (see below for derivation).

### Age Distribution derivation

In a growing population with the doubling time *τ* (approximately equal to the cell cycle time), the number of cells are given approximately given by

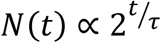

The number of cells of age *a* at time *t* is given by the number of cells of age 0 at time *t* − *a*. Therefore, the distribution of ages *ρρ*(*a*)defined by the number of cells of age *a* divided by total number of cells is given by

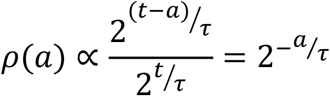

This argument holds for *a* < *τ* but at age *τ*, cells divide and therefore *ρρ*(*a*)= 0 for *a* > *τ*. The proportionality constant can be determined from the normalization

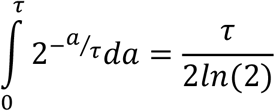

The age distribution is therefore given by

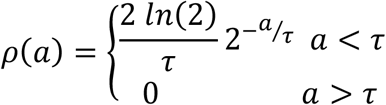

### Averaging over Populations with Heterogeneous Cell-Cycle Times

We can extend these results by considering a population of cells with cell-cycle times *τ*_*c*_ drawn from a distribution *P*(*τ*_*c*_). In this case the population doubling time *τ*_*p*_ would satisfy the following integral equation

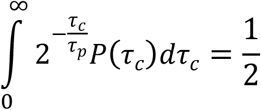

and the age distribution is given by

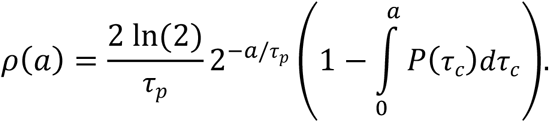

The derivations of these relationships are cited here^70,71^. The logic behind the derivation of the age distribution is very similar to the previous case, except now, instead of cutting the distributions abruptly at some cell-cycle time, it is multiplied by the probability that a cell is not divided until age *a*, which reflects the fact that different cells can divide at different ages.

Now following the previous section, we can find the population averge value of *A* as following

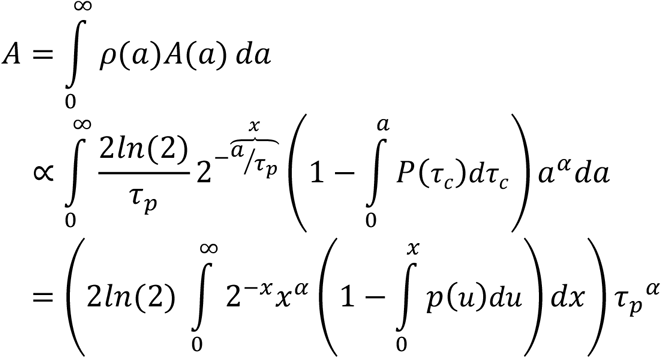

Where *p*(*u*)= *τ*_*p*_ *P*(*τ*_*c*_)is the distribution of cell cycle times rescaled by the population doubling time. If this distribution is similar across different samples (i.e. the relative variability in doubling times are the same across samples), we are essentially done: we have shown that *A* ∝ *τ*_*p*_^*α*^and similarly, *B* ∝ *τ p*^*β*^, and therefore, *B* would scale with *A* as 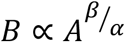.

What if *p*(*u*)is different across samples? Let us approximate the cell cycle time distribution by a gamma distribution of mean *μ* and standard deviation *σ*. Note that narrow gamma distributions are approximately Gaussian by central limit theorem, but they have the advantage of having their support on positive reals. For gamma distributed *p*(*u*), one can show that *A* is approximately proportional to

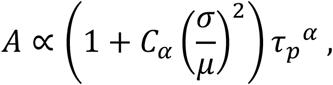

where *C*_*α*_ is an *α*-dependent constant of order 1 (*C*_1_ ≈ 0.7 and *C*_2_ ≈ 1). As long as the vari-ance of (*σ*/*μ*)^2^ across the samples is significantly smaller than the relative variability in population doubling times, the derived scaling of 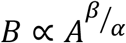 holds. Note that this analysis is done for narrow distributions of cell-cycle times, which already implies (*σ*/*μ*)^2^ ≪ 1 which in turn implies that Var[(*σ*/*μ*)^2^] ≪ 1.

## SUPPLEMENTARY DATA

**Table S1:**
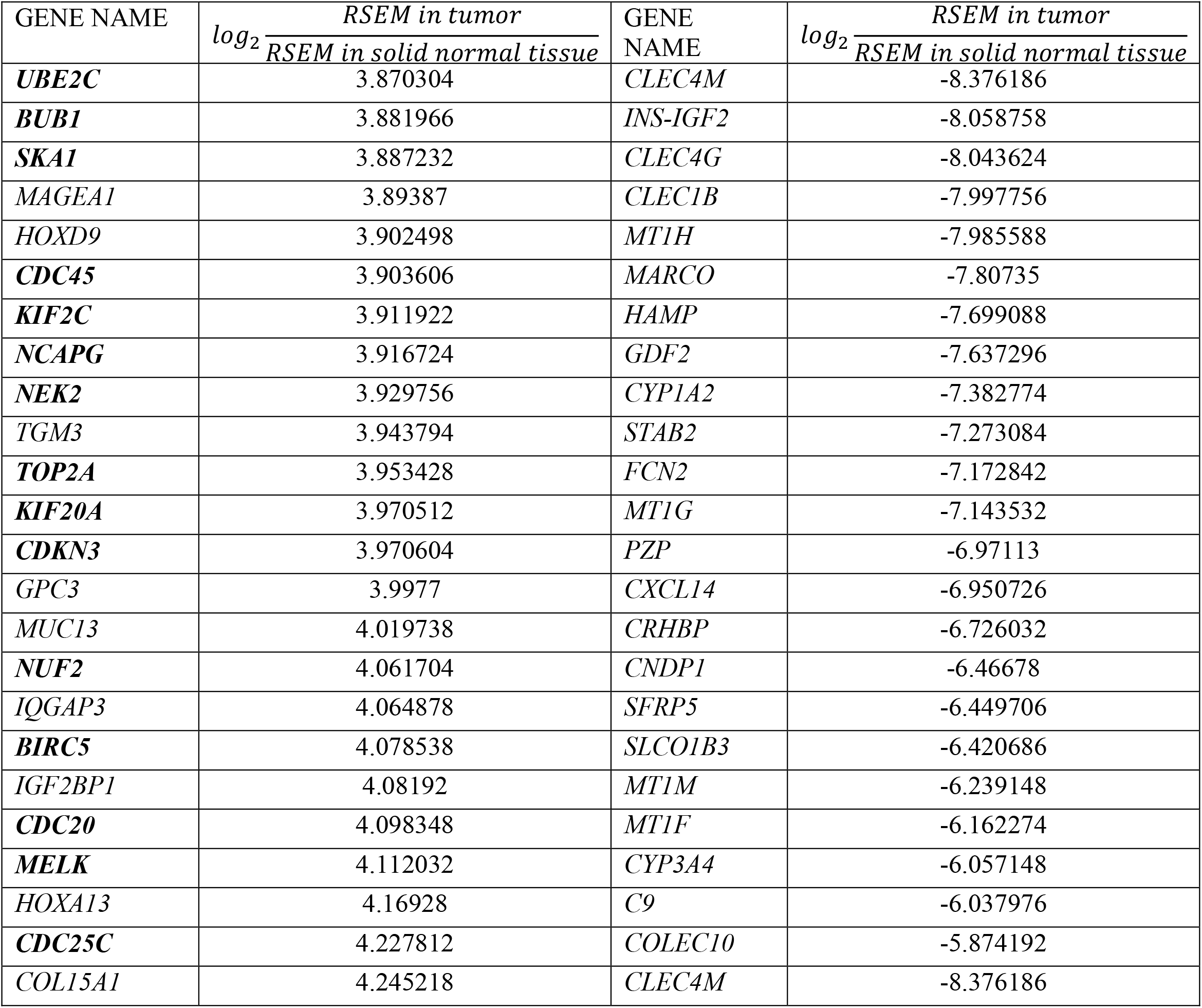
24 genes that have the most increase and most decrease in expression in tumor tissue compared to normal from a subset of 50 patients from the TCGA Liver Cancer dataset. Genes in bold scale strongly with *LMNB1* mRNA expression.

## Supplementary Figure Legends

**Fig. S1.**
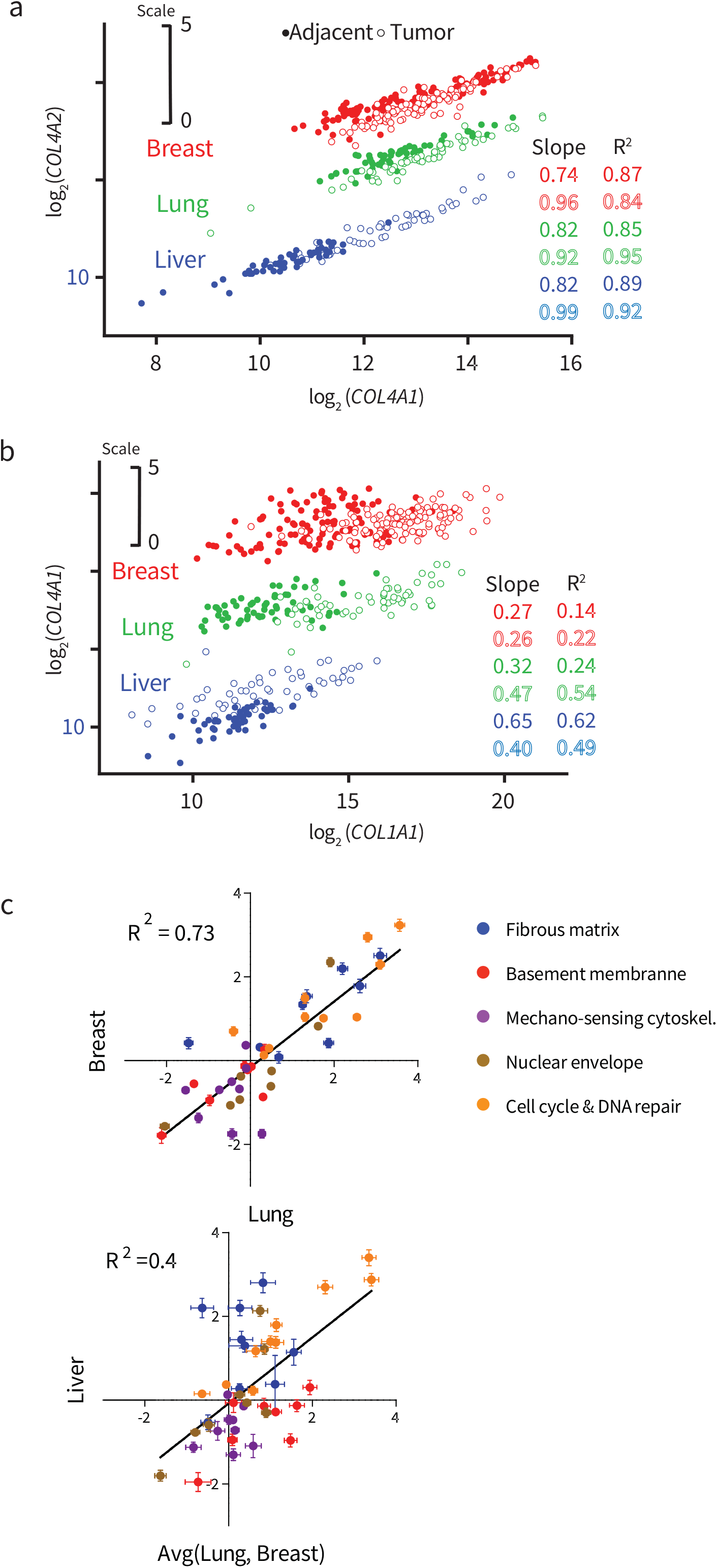
**a)** Plots of log_2_(*COL4A1*) vs log_2_(*COL4A2*) should show an excellent correlation as subunits of the type IV collagen, which are also present very close to each other on the same chromosome and have a shared promotor. Tumor (open circle) and normal (closed circle) tissue data points are plotted individually for Liver (blue), Lung Adenocarcinoma (green) and Breast (red) Cancer. All three cancer datasets show excellent R^2^ (0.84 to 0.95). Tumor data points for only Liver cancer show higher *COL4A1* and *COL4A2* values. (Note that for better visibility, all Lung Adenocarcinoma data points were moved up by 3 units and Breast Cancer by 6 units on the y-axis.) **b)** Plots of log_2_(*COL1A1*) vs log_2_(*COL4A1*) show poor R^2^ (0.14 to 0.62) for most tissue types implying that the basement membrane ECM is regulated differently than fibrous ECM. (Note that for better visibility, all Lung Adenocarcinoma data points were moved up by 5 units and Breast Cancer by 10 units on the y-axis.) **c)** Plotting the fold change in mRNA for Lung vs Breast Cancer shows that genes show similar trends of up/downregulation (R^2^ = 0.73). However, plotting the avg(Lung, Breast) vs Liver Cancer shows a weak correlation (R^2^ = 0.4)

**Fig. S2.**
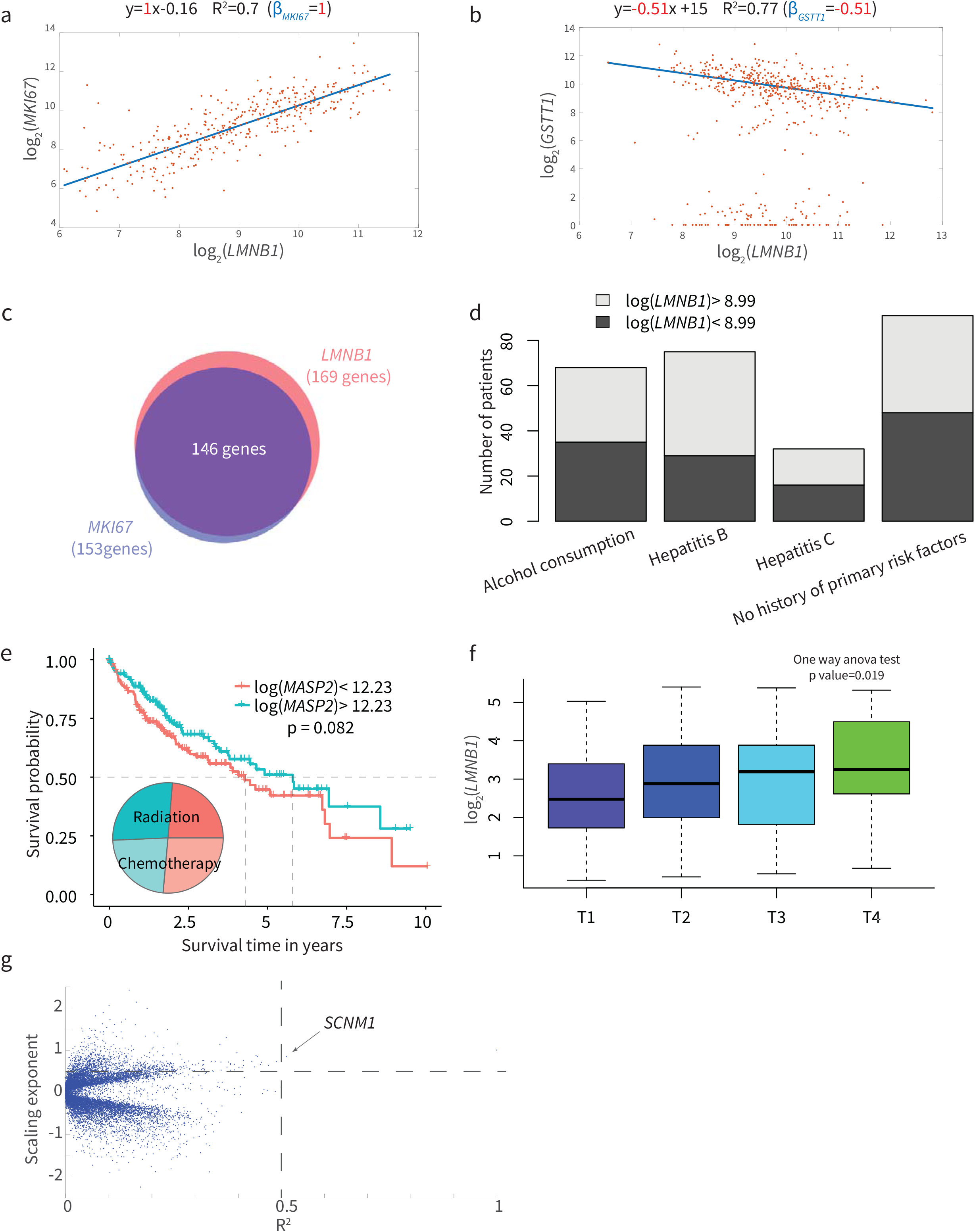
**a)**Plot of log_2_(*LMNB1)* vs log_2_(*MKI67)* in primary Liver Cancer tumor (n=371 patients). **b)** Scatterplot of the only gene that shows a negative power law relation versus *LMNB1* with a good R^2^ = 0.77 in primary Lung Adenocacinoma (n=515 patients), *GSTT1*. **c)** 153 genes are parsed out with slope > 0.5 and R^2^ > 0.5 in the sideways-volcano plot for *MKI67*, 146 of which are identical to genes that scale with *LMNB1*. **d)** Plotting the distribution of primary risk factors including alcohol consumption, hepatitis B and hepatitis C shows no difference between high and low *LMNB1* mRNA expressing patients. **e)** Kaplan-Meier plot for *MASP2* shows that higher expression of *MASP2* increases median survival by ∼2 yrs. **f)** Box and whisker plot of *LMNB1* mRNA stratified on the basis of stage of cancer progression (T1-T4). Increase in median *LMNB1* expression occurs with progression of tumors, with a significant p-value (one-way ANOVA). **g)** Sideways-volcano plot for *LMNA* in primary tumors of Liver Cancer (n=371) shows only one gene with high scaling coefficient (>0.5) and R^2^ > 0.5, which supports the notion that the two lamin isoforms serve distinct functions and reveals *LMNA* does not directly scale with cell proliferation genes.

**Fig. S3.**
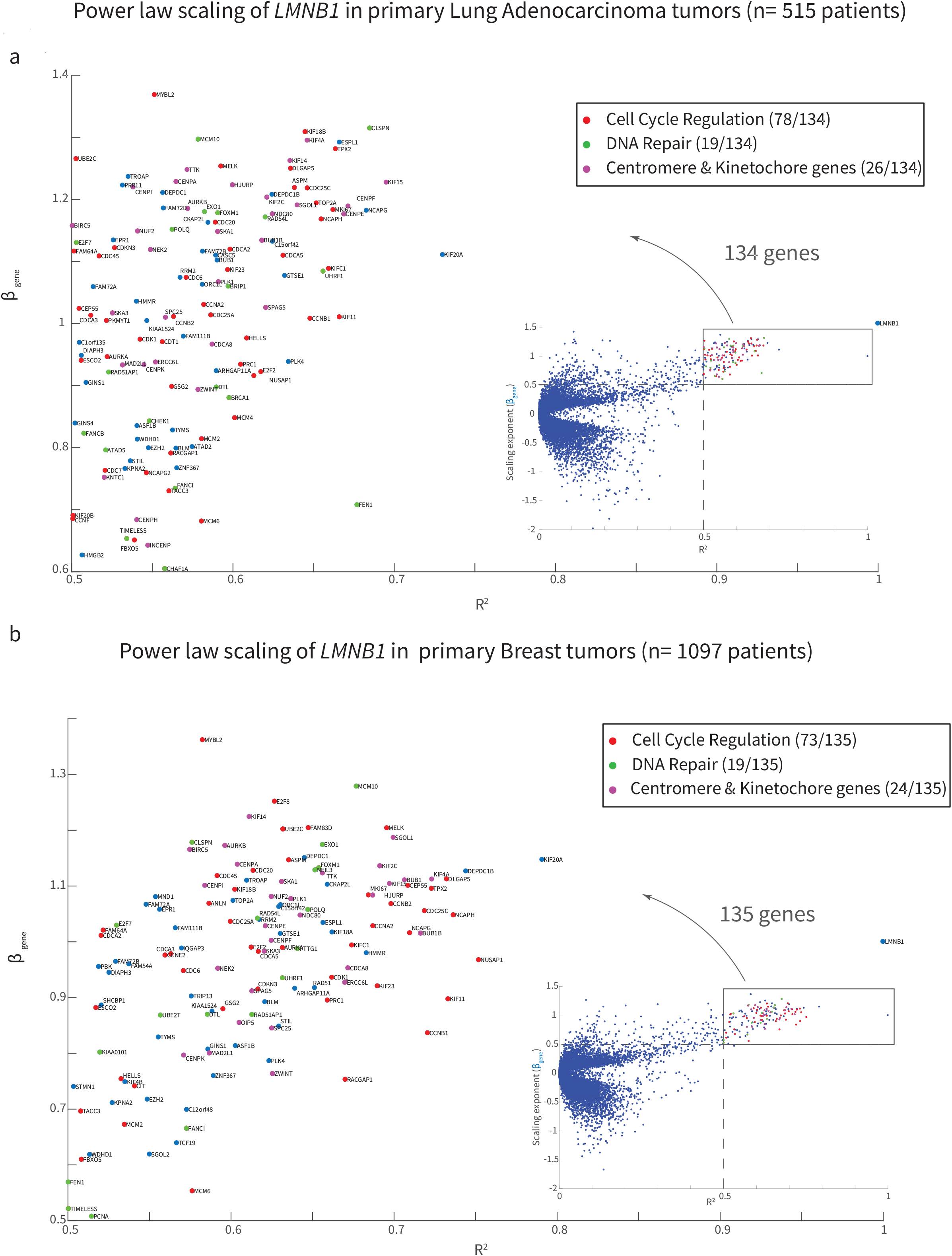
**a)** Power law scaling of *LMNB1* in primary Lung Adenocarcinoma tumors (n=515 patients) similar to Fig 2d shows enrichment of the same annotations (cell cycle regulation, DNA repair, and kinetochore and centromere genes) in the 135 genes with high scaling coefficient *β*_*gene*_ > 0.5 and R^2^ > 0.5; 105 out the 135 genes overlap with the corresponding genes in Liver and Breast Cancer **b)** Similar plot for Power law scaling of *LMNB1* in primary Breast tumors (n=1,097 patients)

**Fig. S4.**
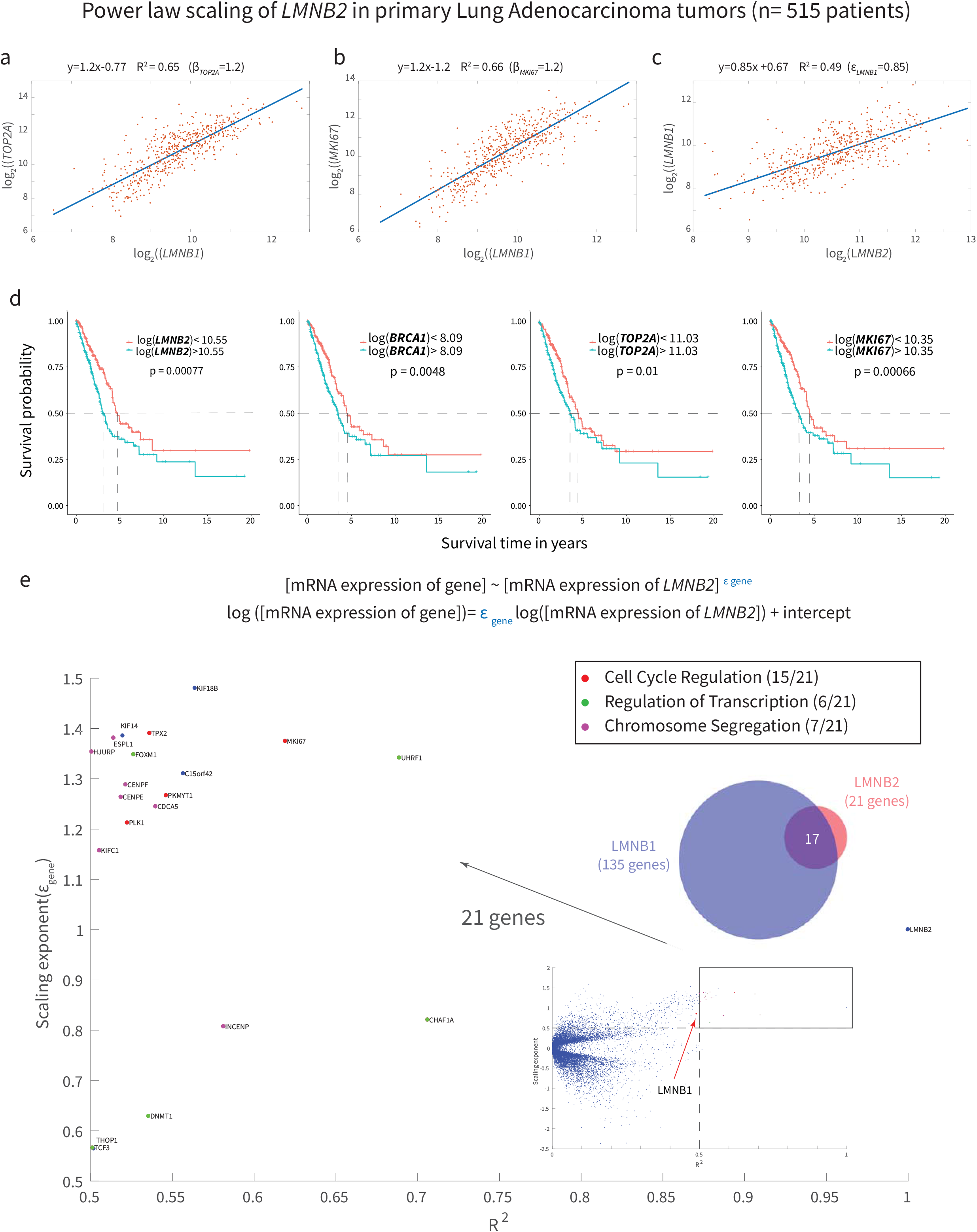
**a-b)** Plot of log_2_(*LMNB1*) vs a) log_2_(*TOP2A*) b) log_2_(*MKI67*) in primary lung adenocarcinoma (n=515 patients) **c)** Plot of log_2_(*LMNB2*) vs log_2_(*LMNB1*) in primary lung adenocarcinoma (n=515 patients) **d)** Kaplan-Meier plot for *LMNB2* shows decrease in survival with increase in expression of *LMNB2* (*p*-value = 0.00077). Similar Kaplan Meier plots for other genes that scale with *LMNB1* (*BRCA1, TOP2A, MKI67*) also show a decrease in survival probability with higher expression. **e)** Genes that scale with *LMNB2*. Parsing genes that show a high scaling coefficient ε_*gene*_ > 0.5 and R^2^>0.5 shows an enrichment of Cell cycle regulation (red), Transcription regulation (green) and Chromosome segregation genes (purple). Top right inset: Venn diagram for genes parsed out for *LMNB2* and *LMNB1* (Fig. S3) shows that 17 out of 21 genes for *LMNB2* are identical to those of *LMNB1*.

**Fig. S5.**
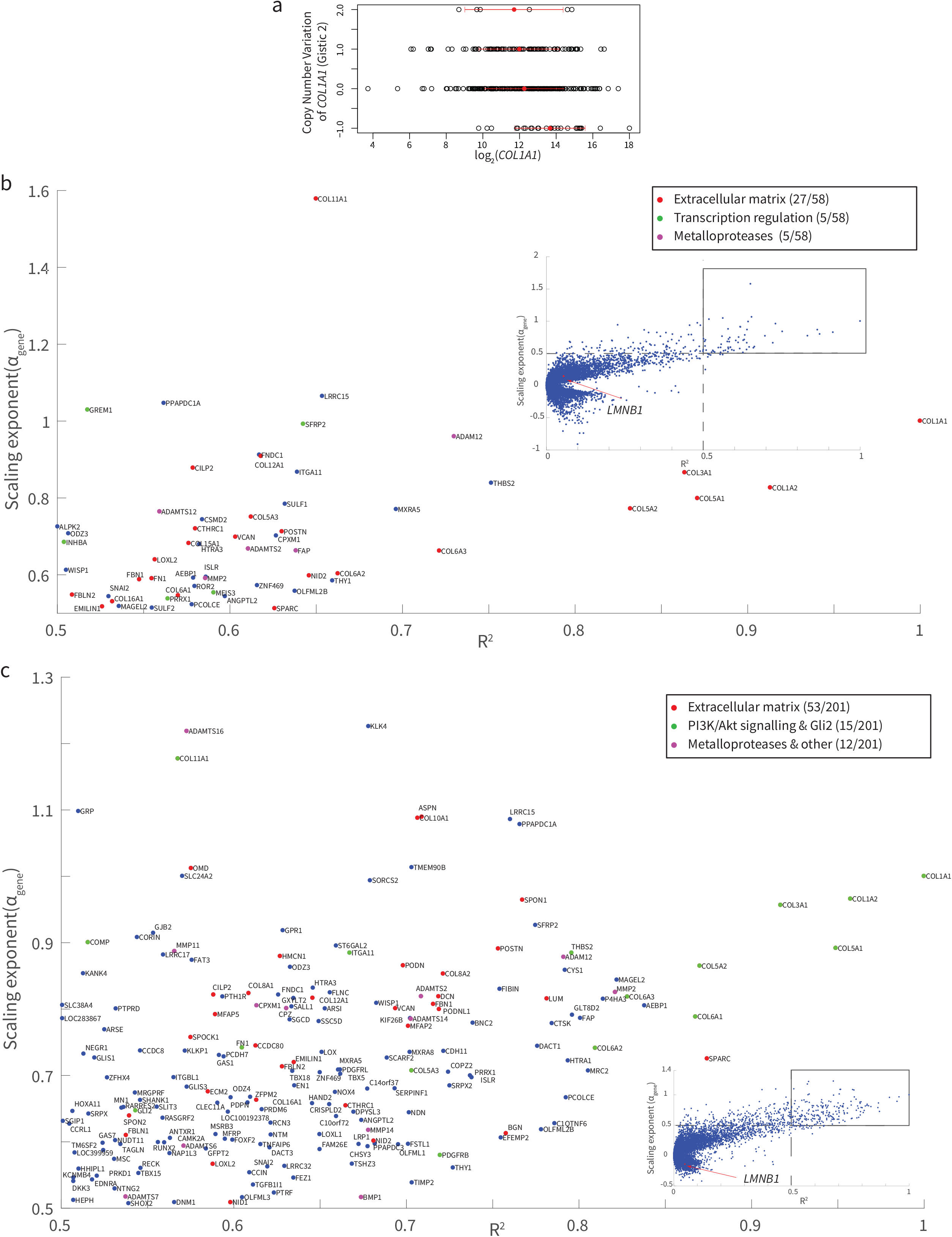
***a)*** *COL1A1* mRNA for primary Liver Cancer tumor (n=371) vs the corresponding threshold Gistic 2 copy number variation data for the same patient. The Gistic2 values have a thresholding of -2, -1,0,1,2, representing homozygous deletion, single copy deletion, diploid normal copy, low-level copy number amplification, or high-level copy number amplification respectively. The red points indicate the mean mRNA expression values. **b)** Power law scaling of *COL1A1* in primary Lung Adenocarcinoma tumors (n=515 patients) similar to **Fig. 3d** shows enrichment of similar annotations (extracellular matrix, transcription regulation, metalloproteases) in the 58 genes with high scaling coefficient *α*_*gene*_ > 0.5 and R^2^ > 0.5. **c)** Similar plot for *COL1A1* in primary Breast tumors (n=1,097 patients).

**Fig. S6.**
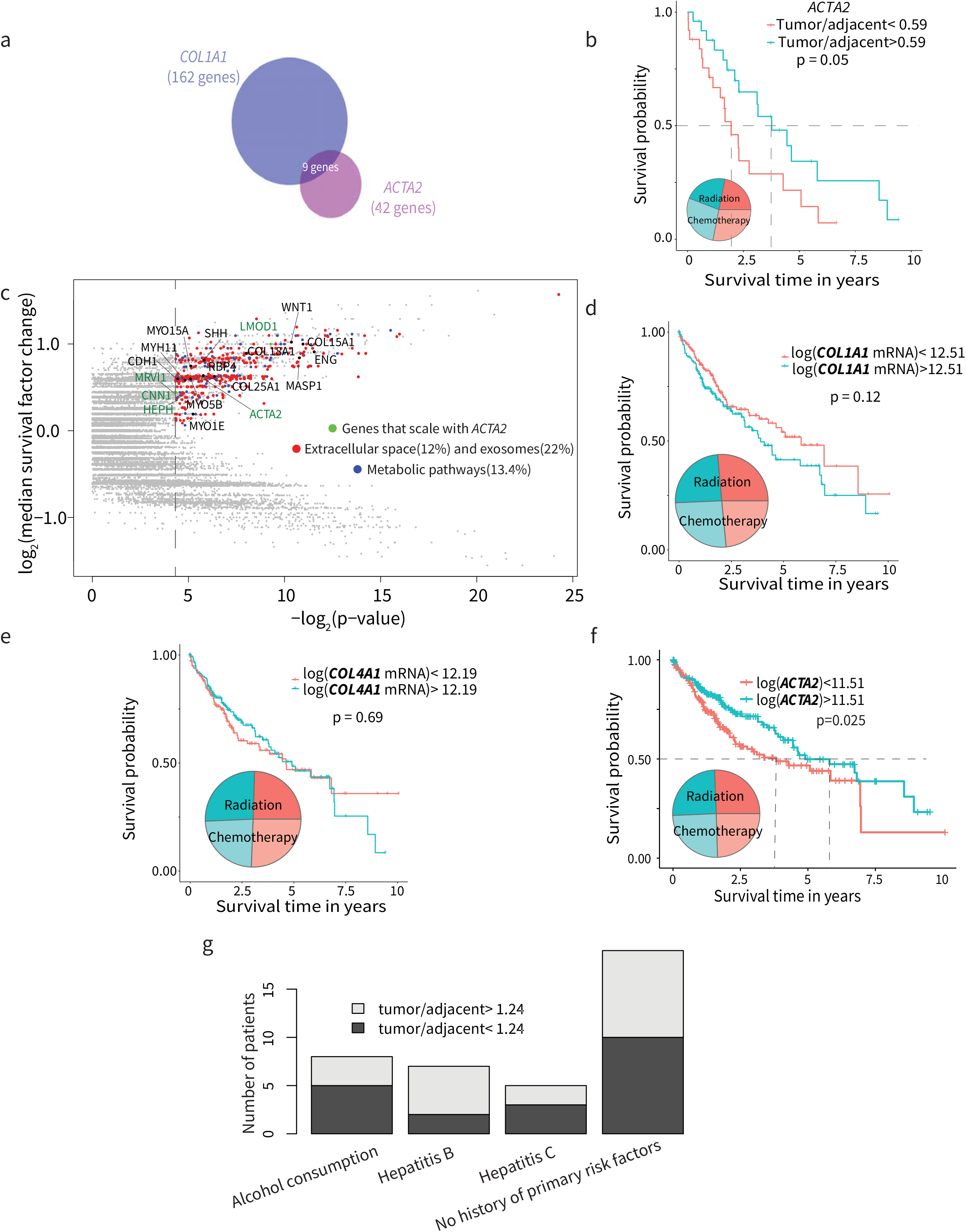
**a)** Sideways-Volcano plots for *COL1A1* and *ACTA2* for 371 Liver primary tumors gives 9 overlapping genes with slope > 0.5 and R^2^ > 0.5. **b)** Kaplan-Meier plot shows patients with a higher upregulation of *ACTA2* expression in tumors compared to patient matched adjacent uninvolved tissue have significantly higher survival probability (n=50 patients). **c)** Gene Ontology enrichment analysis of pro-survival genes in the Whole Genome Survival Plot. **d)** *COL1A1* mRNA levels alone are not a good indicator of survival probability. **e)** Basement membrane collagen *COL4A1* likewise does not predict survival likelihood. **f)** Dividing the range of marker for activated fibroblasts-*ACTA2* mRNA levels (in primary tumor of 371 patients) into two groups, one transcribing higher *ACTA2* mRNA(greater than the median) and the other lower, and plotting a Kaplan Meier(KM) using the phenotype data available for time of death, we find that patients translating higher *ACTA2* mRNA in the tumor tissue possess a significantly lower risk of death than ones with lower *ACTA2* (median survival (number of years after which survival probability = 0.5) reduced by ∼2 years). **g)**Plotting the distribution of primary risk factors including alcohol consumption, hepatitis B and hepatitis C shows no difference between high and low “uninvolved-tissue normalized” tumor level of *COL1A1*.

**Fig. S7.**
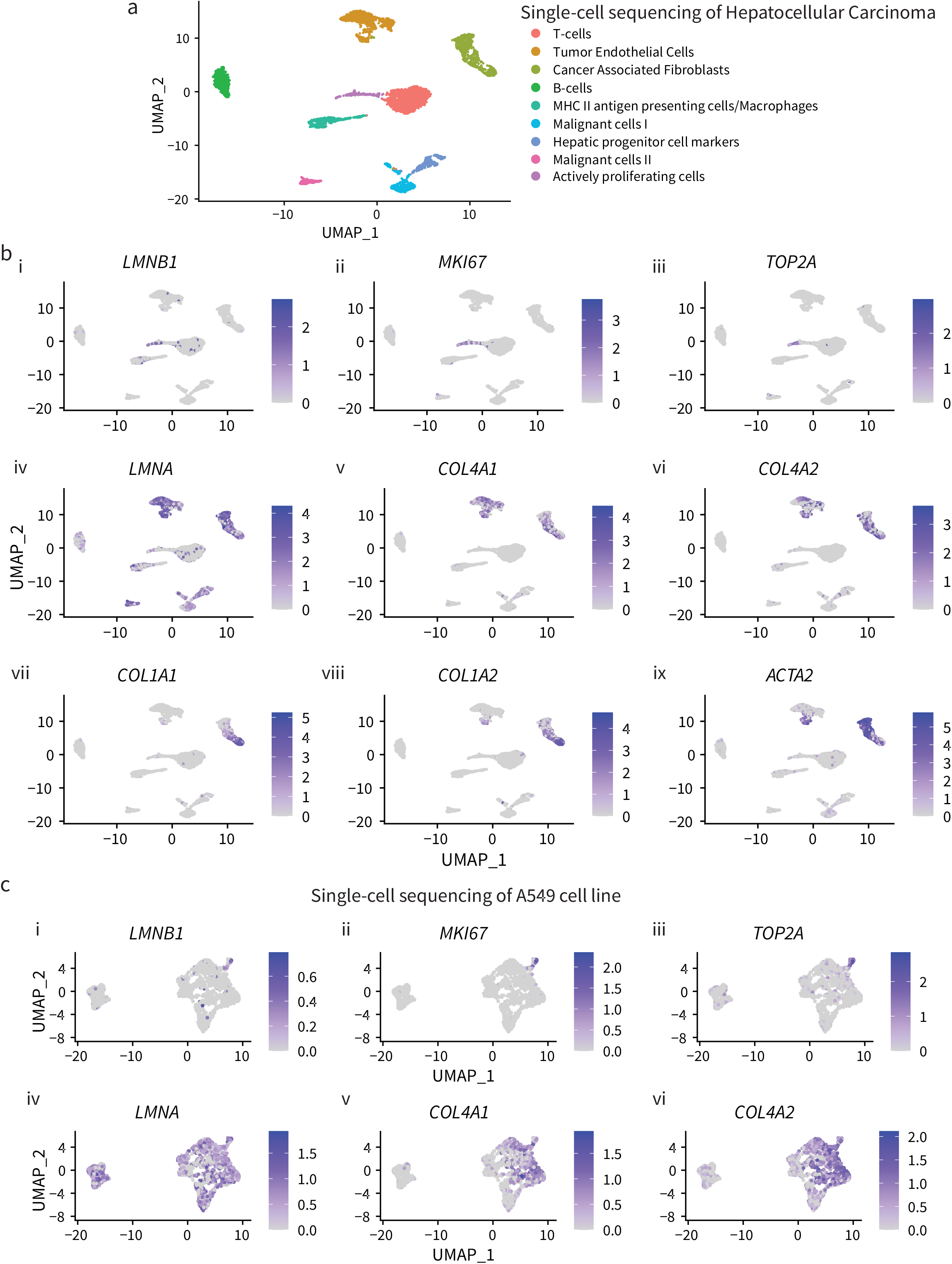
**a)** Dimensionally reduced projection (UMAP) of single-cell mRNA sequencing of HCC tumors depicting clusters based on sequencing profiles labeled with the corresponding cell-type phenotype **b)** UMAP from (a) labeled with expression levels of *LMNB1, MKI67, TOP2A, LMNA, COL4A1, COL4A2, COL1A1, COL1A2, ACTA2* in each cell of the clusters **c)** Dimensionally reduced projection (UMAP) of single-cell mRNA sequencing of A549 cells labeled with expression levels of *LMNB1, MKI67, TOP2A, LMNA, COL4A1, COL4A2*

**Fig. S8.**
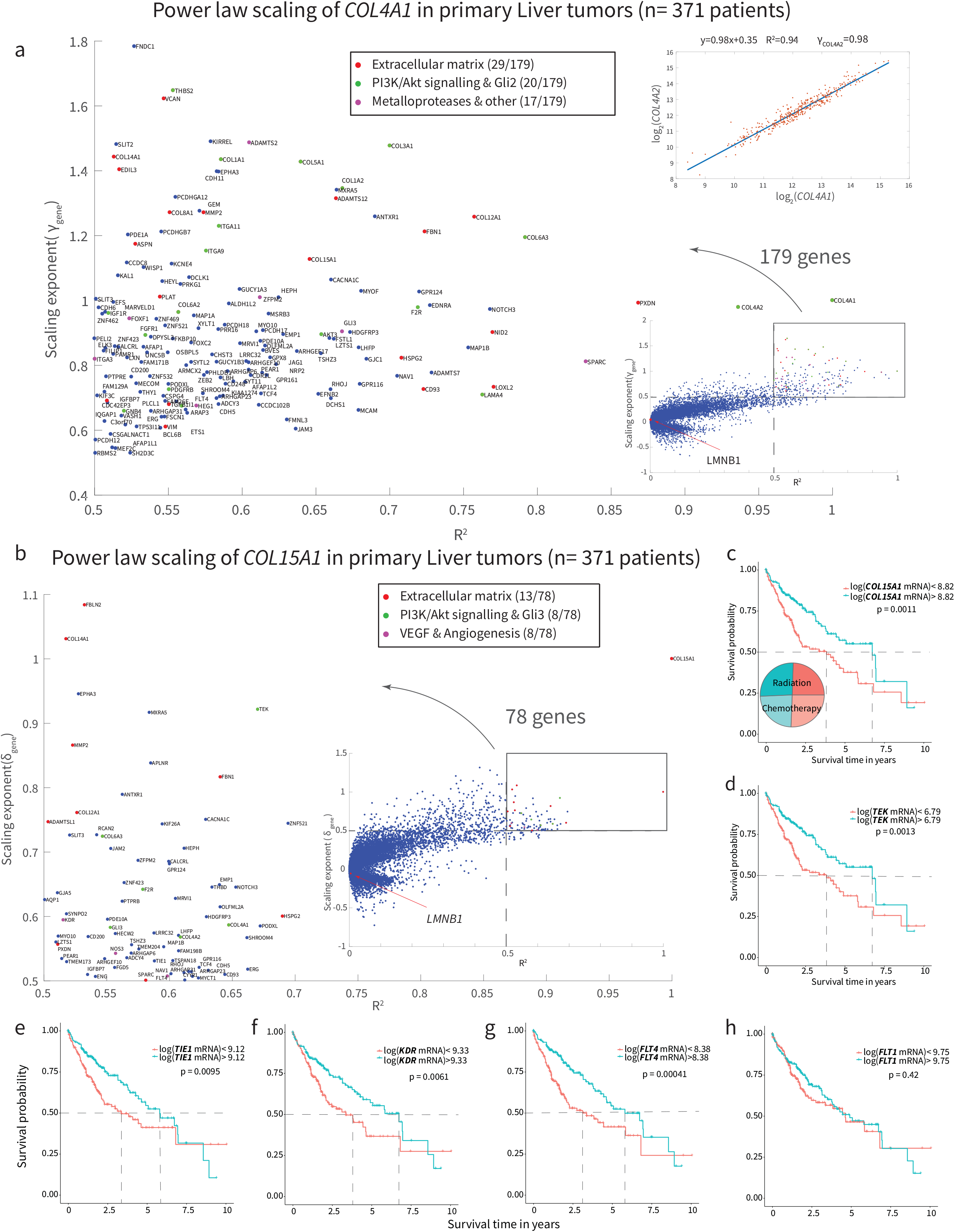
**a)** Power law scaling of *COL4A1* in primary Liver tumors (n=371 patients) similar to **Fig.3d** shows enrichment of the same annotations (extracellular matrix, *PI3K/Akt* signaling & *GLI2* and metalloproteases & other ECM remodeling enzymes) in the 163 genes with high scaling coefficient *γ*_*gene*_ >0.5 and R^2^ > 0.5. *γ*_*COL*4*AA*2_ ∼1 and R^2^ = 0.93 also validates the dataset as *COL4A1* and *COL4A2* are subunits of the type IV collagen heterotrimer. **b)** Similar plot for *COL15A1* shows enrichment of the extracellular matrix (red), *PI3K/Akt* signaling & *GLI2* (green) and VEGF & angiogenesis (purple) annotations in the 78 genes with high scaling coefficient δ_*gene*_ > 0.5 and R^2^ > 0.5. **c)** Patients with higher *COL15A1* mRNA in the tumor tissue (greater than the median) possess a significantly higher probability of survival than ones with lower *COL15A1*. **d-g)** Similar Kaplan-Meier plot for other genes that scale with *COL15A1* (*TEK, TIE1, FLT4, KDR*) show similar increases in survival probability with higher expression. **h)** Kaplan-Meier plot for *FLT1*, which is also a receptor tyrosine kinase of VEGF but does not have a good power law fit with *COL15A1*, does not show any significant increase in survival probability with higher expression of the gene.

**Fig.S9.**
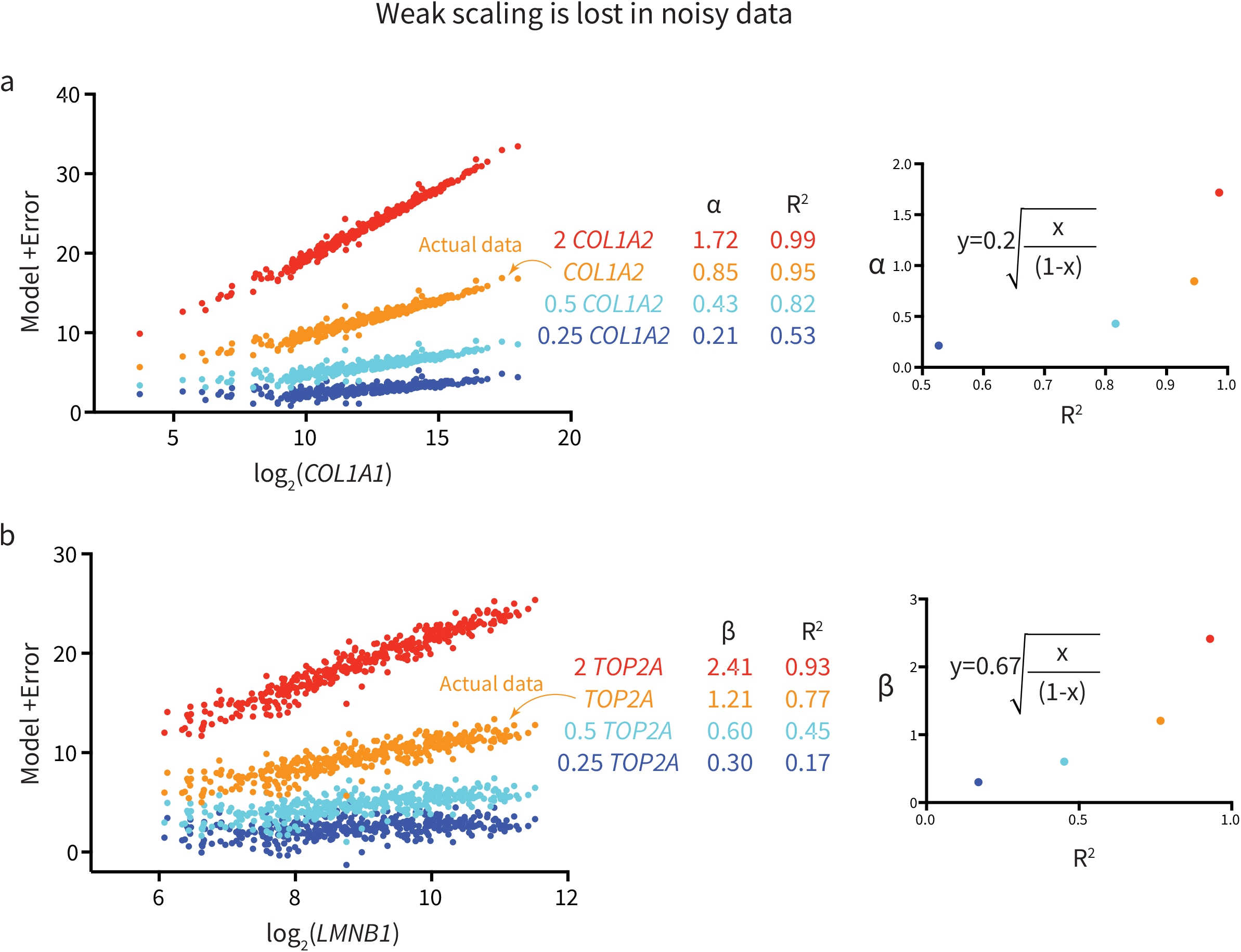
Weak scaling can be lost in noisy data. **a)** Multiplying the fit of *log*_2_([*mRNA expression of COL*1*A*1])vs *log*_2_([*mRNA expression of COL*1*A*2])(orange) by 2 (red), 0.5 (cyan) and 0.25 (blue) and adding back the residuals of the fit with a RMSE of 0.44, gives lower R^2^ for lower slope. Right plot: Fitting the different slopes vs R2 for the same RMSE. **b)** Similarly for model fit obtained from *log*_2_([*mRNA expression of CCBAB*1])vs *log*_2_([*mRNA expression of TOP*2*A*])with a higher RMSE of 0.82.

## References

1. Levental, K. R. et al. Matrix crosslinking forces tumor progression by enhancing integrin signaling. Cell 139, 891–906 (2009).

2. Pinter, M., Trauner, M., Peck-Radosavljevic, M. & Sieghart, W. Cancer and liver cirrhosis: implications on prognosis and management. ESMO open 1, e000042 (2016).

3. Saito, A., Horie, M., Micke, P. & Nagase, T. The role of TGF-β signaling in lung cancer associated with idiopathic pulmonary fibrosis. International Journal of Molecular Sciences 19, (2018).

4. Harada, T. et al. Nuclear lamin stiffness is a barrier to 3D migration, but softness can limit survival. J. Cell Biol. 204, 669–682 (2014).

5. Dahl, K. N., Engler, A. J., Pajerowski, J. D. & Discher, D. E. Power-law rheology of isolated nuclei with deformation mapping of nuclear substructures. Biophys. J. 89, 2855–2864 (2005).

6. Pajerowski, J. D., Dahl, K. N., Zhong, F. L., Sammak, P. J. & Discher, D. E. Physical plasticity of the nucleus in stem cell differentiation. PNAS 104, 15619–15624 (2007).

7. Irianto, J. et al. Nuclear constriction segregates mobile nuclear proteins away from chromatin. Mol. Biol. Cell 27, 4011–4020 (2016).

8. Shimi, T. et al. The role of nuclear lamin B1 in cell proliferation and senescence. Genes Dev. 25, 2579–93 (2011).

9. Cho, S. et al. Mechanosensing by the Lamina Protects against Nuclear Rupture, DNA Damage, and Cell-Cycle Arrest. Dev. Cell 49, 920-935.e5 (2019).

10. Klein, E. A. et al. Cell-cycle control by physiological matrix elasticity and in vivo tissue stiffening. Curr. Biol. 19, 1511–8 (2009).

11. Meng, Z. et al. RAP2 mediates mechanoresponses of the Hippo pathway. Nature 560, 655–660 (2018).

12. Swift, J. et al. Nuclear Lamin-A Scales with Tissue Stiffness and Enhances Matrix-Directed Differentiation. Science (80-.). 341, 1240104–1240104 (2013).

13. Yang, Y., Leone, L. M. & Kaufman, L. J. Elastic Moduli of Collagen Gels Can Be Predicted from Two-Dimensional Confocal Microscopy. Biophys. J. 97, 2051–2060 (2009).

14. Kleiber, M. Body size and metabolic rate. Physiol. Rev. 27, 511–541 (1947).

15. Thommen, A. et al. Body size-dependent energy storage causes Kleiber’s law scaling of the metabolic rate in planarians. Elife 8, (2019).

16. Swift, J. et al. Nuclear Lamin-A Scales with Tissue Stiffness and Enhances Matrix-Directed Differentiation. doi:10.1126/science.1243643

17. Cho, S., Irianto, J. & Discher, D. E. Mechanosensing by the nucleus: From pathways to scaling relationships. J. Cell Biol. 216, 305–315 (2017).

18. Qayyum, A. et al. Immunotherapy response evaluation with magnetic resonance elastography (MRE) in advanced HCC. J. Immunother. Cancer 7, 329 (2019).

19. Irianto, J., Pfeifer, C. R., Ivanovska, I. L., Swift, J. & Discher, D. E. Nuclear Lamins in Cancer. Cell. Mol. Bioeng. 9, 258–267 (2016).

20. Sun, S., Xu, M. Z., Poon, R. T., Day, P. J. & Luk, J. M. Circulating Lamin B1 (LMNB1) Biomarker Detects Early Stages of Liver Cancer in Patients. J. Proteome Res. 9, 70–78 (2010).

21. Abdelghany, A. M. et al. Using Lamin B1 mRNA for the early diagnosis of hepatocellular carcinoma: a cross-sectional diagnostic accuracy study. F1000Research 7, 1339 (2018).

22. Vergnes, L., Péterfy, M., Bergo, M. O., Young, S. G. & Reue, K. Lamin B1 is required for mouse development and nuclear integrity. Proc. Natl. Acad. Sci. U. S. A. 101, 10428–10433 (2004).

23. Coffinier, C. et al. Deficiencies in lamin B1 and lamin B2 cause neurodevelopmental defects and distinct nuclear shape abnormalities in neurons. Mol. Biol. Cell 22, 4683–93 (2011).

24. Dou, Z. et al. Autophagy mediates degradation of nuclear lamina. Nature 527, 105–109 (2015).

25. de la Rosa, J. et al. Prelamin A causes progeria through cell-extrinsic mechanisms and prevents cancer invasion. Nat. Commun. 4, 2268 (2013).

26. Hernandez, L. et al. Functional Coupling between the Extracellular Matrix and Nuclear Lamina by Wnt Signaling in Progeria. Dev. Cell 19, 413–425 (2010).

27. Buxboim, A. et al. Matrix elasticity regulates lamin-A,C phosphorylation and turnover with feedback to actomyosin. Curr. Biol. 24, 1909–17 (2014).

28. Dasika, G. K. et al. DNA damage-induced cell cycle checkpoints and DNA strand break repair in development and tumorigenesis. Oncogene 18, 7883–7899 (1999).

29. Kaspi, E. et al. Low lamin A expression in lung adenocarcinoma cells from pleural effusions is a pejorative factor associated with high number of metastatic sites and poor Performance status. PLoS One 12, e0183136 (2017).

30. Capo-chichi, C. D. et al. Loss of A-type lamin expression compromises nuclear envelope integrity in breast cancer. Chin. J. Cancer 30, 415–25 (2011).

31. Harada, T. et al. Nuclear lamin stiffness is a barrier to 3D migration, but softness can limit survival. J Cell Biol 204, 669–682 (2014).

32. Discher, D. E., Janmey, P. & Wang, Y.-L. Tissue Cells Feel and Respond to the Stiffness of Their Substrate. Science (80-.). 310, 1139–1143 (2005).

33. Ho, C. Y., Jaalouk, D. E., Vartiainen, M. K. & Lammerding, J. Lamin A/C and emerin regulate MKL1–SRF activity by modulating actin dynamics. Nature 497, 507–511 (2013).

34. Dupont, S. et al. Role of YAP/TAZ in mechanotransduction. Nature 474, 179–183 (2011).

35. Miralles, F., Posern, G., Zaromytidou, A.-I. & Treisman, R. Actin Dynamics Control SRF Activity by Regulation of Its Coactivator MAL. Cell 113, 329–342 (2003).

36. Palozola, K. C. et al. Mitotic transcription and waves of gene reactivation during mitotic exit. Science (80-.). 358, 119–122 (2017).

37. Mermel, C. H. et al. GISTIC2.0 facilitates sensitive and confident localization of the targets of focal somatic copy-number alteration in human cancers. Genome Biol. 12, R41 (2011).

38. Bradley, M. W., Aiello, K. A., Ponnapalli, S. P., Hanson, H. A. & Alter, O. GSVD-and tensor GSVD-uncovered patterns of DNA copy-number alterations predict adenocarcinomas survival in general and in response to platinum. APL Bioeng. 3, 036104 (2019).

39. Sankaranarayanan, P., Schomay, T. E., Aiello, K. A. & Alter, O. Tensor GSVD of Patient-and Platform-Matched Tumor and Normal DNA Copy-Number Profiles Uncovers Chromosome Arm-Wide Patterns of Tumor-Exclusive Platform-Consistent Alterations Encoding for Cell Transformation and Predicting Ovarian Cancer Survival. PLoS One 10, e0121396 (2015).

40. Aiello, K. A. & Alter, O. Platform-Independent Genome-Wide Pattern of DNA Copy-Number Alterations Predicting Astrocytoma Survival and Response to Treatment Revealed by the GSVD Formulated as a Comparative Spectral Decomposition. PLoS One 11, e0164546 (2016).

41. Pfau, S. J. & Amon, A. Chromosomal instability and aneuploidy in cancer: from yeast to man. EMBO Rep. 13, 515–27 (2012).

42. Han, L. et al. Lamin B2 Levels Regulate Polyploidization of Cardiomyocyte Nuclei and Myocardial Regeneration Article Lamin B2 Levels Regulate Polyploidization of Cardiomyocyte Nuclei and Myocardial Regeneration. Dev. Cell1–18 (2020). doi:10.1016/j.devcel.2020.01.030

43. Pfeifer, C. R. et al. Constricted migration increases DNA damage and independently represses cell cycle. Mol. Biol. Cell 29, 1948–1962 (2018).

44. Gerace, L. & Blobel, G. The nuclear envelope lamina is reversibly depolymerized during mitosis. Cell 19, 277–87 (1980).

45. Gelse, K., Pöschl, E. & Aigner, T. Collagens--structure, function, and biosynthesis. Adv. Drug Deliv. Rev. 55, 1531–46 (2003).

46. Hofheinz, R.-D. et al. Stromal Antigen Targeting by a Humanised Monoclonal Antibody: An Early Phase II Trial of Sibrotuzumab in Patients with Metastatic Colorectal Cancer. Oncol. Res. Treat. 26, 44–48 (2003).

47. Wells, R. G. The Role of Matrix Stiffness in Hepatic Stellate Cell Activation and Liver Fibrosis. J. Clin. Gastroenterol. 39, S158–S161 (2005).

48. Ma, L. et al. Tumor Cell Biodiversity Drives Microenvironmental Reprogramming in Liver Cancer. Cancer Cell 36, 418–430.e6 (2019).

49. Dittmer, T. A. & Misteli, T. The lamin protein family. Genome Biol. 12, 222 (2011).

50. Peter, A. & Stick, R. Evolution of the lamin protein family: What introns can tell. Nucleus 3, 44–59 (2012).

51. Lopez-Soler, R. I., Moir, R. D., Spann, T. P., Stick, R. & Goldman, R. D. A role for nuclear lamins in nuclear envelope assembly. J. Cell Biol. 154, 61–70 (2001).

52. Moir, R. D., Yoon, M., Khuon, S. & Goldman, R. D. Nuclear Lamins A and B1: Different Pathways of Assembly during Nuclear Envelope Formation in Living Cells. The Journal of Cell Biology 151, (2000).

53. Harborth, J., Elbashir, S. M., Bechert, K., Tuschl, T. & Weber, K. Identification of essential genes in cultured mammalian cells using small interfering RNAs. J. Cell Sci. 114, 4557–65 (2001).

54. Moir, R. D., Spann, T. P., Herrmann, H. & Goldman, R. D. isruption of Nuclear Lamin Organization Blocks the Elongation Phase of DNA Replication. The Journal of Cell Biology 149, (2000).

55. Moir, R. D., Montag-Lowy, M. & Goldman, R. D. Dynamic properties of nuclear lamins: lamin B is associated with sites of DNA replication. J. Cell Biol. 125, 1201–12 (1994).

56. Omenetti, A., Choi, S., Michelotti, G. & Diehl, A. M. Hedgehog signaling in the liver. J. Hepatol. 54, 366–73 (2011).

57. Lemoinne, S. et al. Portal myofibroblasts promote vascular remodeling underlying cirrhosis formation through the release of microparticles. Hepatology 61, 1041–1055 (2015).

58. Özdemir, B. C. et al. Depletion of Carcinoma-Associated Fibroblasts and Fibrosis Induces Immunosuppression and Accelerates Pancreas Cancer with Reduced Survival. Cancer Cell 25, 719–734 (2014).

59. Edenberg, H. J. The genetics of alcohol metabolism: role of alcohol dehydrogenase and aldehyde dehydrogenase variants. Alcohol Res. Health 30, 5–13 (2007).

60. Crabb, D. W., Matsumoto, M., Chang, D. & You, M. Overview of the role of alcohol dehydrogenase and aldehyde dehydrogenase and their variants in the genesis of alcohol-related pathology. Proc. Nutr. Soc. 63, 49–63 (2004).

61. Leu, J. I.-J., Dumont, P., Hafey, M., Murphy, M. E. & George, D. L. Mitochondrial p53 activates Bak and causes disruption of a Bak–Mcl1 complex. Nat. Cell Biol. 6, 443–450 (2004).

62. Li, L. Y., Luo, X. & Wang, X. Endonuclease G is an apoptotic DNase when released from mitochondria. Nature 412, 95–99 (2001).

63. Du, C., Fang, M., Li, Y., Li, L. & Wang, X. Smac, a Mitochondrial Protein that Promotes Cytochrome c– Dependent Caspase Activation by Eliminating IAP Inhibition. Cell 102, 33–42 (2000).

64. Susin, S. A. et al. Molecular characterization of mitochondrial apoptosis-inducing factor. Nature 397, 441–446 (1999).

65. Chun, Y. S. et al. Significance of pathologic response to preoperative therapy in pancreatic cancer. Ann. Surg. Oncol. 18, 3601–3607 (2011).

66. Motosugi, U. et al. Liver stiffness measured by magnetic resonance elastography as a risk factor for hepatocellular carcinoma: A preliminary case-control study. Eur. Radiol. 23, 156–162 (2013).

67. Batheja, M. et al. Magnetic resonance elastography (MRE) in assessing hepatic fibrosis: performance in a cohort of patients with histological data. Abdom. Imaging 40, 760–765 (2015).

68. Tamaki, N. et al. Risk assessment of hepatocellular carcinoma development by magnetic resonance elastography in chronic hepatitis C patients who achieved sustained virological responses by direct-acting antivirals. J. Viral Hepat. 26, 893–899 (2019).

69. Butler, A., Hoffman, P., Smibert, P., Papalexi, E. & Satija, R. Integrating single-cell transcriptomic data across different conditions, technologies, and species. Nat. Biotechnol. 36, 411–420 (2018).

70. Powell, E. O. Growth Rate and Generation Time of Bacteria, with Special Reference to Continuous Culture. J. Gen. Microbiol. 15, 492–511 (1956).

71. Jafarpour, F. et al. Bridging the Timescales of Single-Cell and Population Dynamics. Phys. Rev. X 8, 021007 (2018).

72. Collisson, E. A. et al. Comprehensive molecular profiling of lung adenocarcinoma: The cancer genome atlas research network. Nature 511, 543–550 (2014).

